# NSC348884 cytotoxicity is not mediated by inhibition of nucleophosmin oligomerization

**DOI:** 10.1101/2020.02.06.936666

**Authors:** Markéta Šašinková, Petr Heřman, Aleš Holoubek, Dita Strachotová, Petra Otevřelová, Dana Grebeňová, Kateřina Kuželová, Barbora Brodská

**Author notes:** Corresponding authors: Barbora Brodská, Department of Proteomics, Institute of Hematology and Blood Transfusion, U Nemocnice 1, 128 20 Prague 2, Czech Republic, Petr Heřman, Faculty of Mathematics and Physics, Institute of Physics, Charles University, Ke Karlovu 5, 121 16 Prague 2, Czech Republic.

## Abstract

Oligomerization of the nucleolar phosphoprotein nucleophosmin (NPM) is mediated by its N-terminal domain. In acute myeloid leukemia, a frequent NPM mutation occurring at the C-terminus causes NPM delocalization to the cytoplasm. Due to formation of NPM heterooligomers, the wild-type NPM as well as many of NPM interaction partners are also delocalized. Proper localization and function of mislocalized proteins in the cells with mutated NPM may be restored by targeting NPM oligomerization. We introduce a reliable set of complementary methods for monitoring NPM oligomerization in both cell lysates and live cells. Using this methodological background we show that a putative inhibitor of NPM oligomerization, NSC348884, does not prevent formation of NPM oligomers in leukemia cells. Instead, we reveal that the observed cytotoxic effect of NSC348884 is associated with changes in cell adhesion signaling.

## Introduction

The human *NPM1* gene is located on the chromosome 5q35 and encodes a 32,6 kDa polypeptide. Nucleophosmin (NPM), encoded by the *NPM1* gene, is a ubiquitously expressed phosphoprotein residing predominantly in the granular component of the nucleolus and dynamically shuttling among nucleoli, the nucleoplasm and the cytoplasm (4, 11, 12, 52). It functions as a chaperone (24, 46) and regulates various cellular processes including the ribosome biogenesis (34, 67, 69), DNA-damage repair (58, 75, 77), centrosome duplication (53, 74), and DNA replication (54, 71, 72). Furthermore, NPM is involved in apoptosis and can modulate p53 stability and activity (11, 43, 75).

NPM overexpression, fusion, or mutation have oncogenic potential and are associated with cancer progression in many types of solid tumors (29) and in hematopoietic malignancies (8, 20, 50, 63, 76). Acute myeloid leukemia (AML) with mutated *NPM1* accounts for about 1/3 of de novo adult AML, and *NPM1* is the most frequently mutated gene in AML with normal karyotype (50-60 % incidence) (18, 20, 21). To date, more than 100 *NPM1* mutation types have been identified in AML, occurring almost exclusively in the last exon (exon 12) of the gene (5). Rare mutations have been detected in the exons 5, 6, 9, and 11 (1, 48, 57). A nucleotide insertion and/or deletion (indel mutations) leads to the frame shift in the region encoding the C-terminus of NPM. All these *NPM1* mutations result in loss of tryptophan residues at positions 288 and/or 290, which form the main part of the nucleolar localization signal (NoLS) ensuring nucleolar localization of the wild-type NPM (NPMwt). Moreover, the indel mutations frequently generate an additional nuclear export sequence (NES), which labels the mutated protein for the nuclear exporter XPO1 and targets NPM into the cytoplasm (22). Both the loss of NoLS and the occurrence of the new NES lead to the accumulation of the NPM mutant (NPMmut) in the cytoplasm (18, 20, 21). Cytoplasmic NPM serves as an immunohistochemical marker with prognostic relevance (14, 19, 26, 39, 70, 73), and is also associated with reduced incidence of some HLA class I alleles, possibly due to anti-leukemia immune response (40, 41). It has been shown that homozygous mutation of *NPM1* causes early embryonic lethality (9, 28). Accordingly, AML patients with *NPM1* mutation are always heterozygous (20, 21). In the absence of an additional genetic aberration, they have better response to intensive chemotherapy. Nevertheless, AML at elderly patients is often associated with other chromosomal and gene abnormalities, in particular with mutations in *DNMT3A, TET2, TP53* or *ASXL1* genes, which have adverse impact on prognosis and overcome the influence of *NPM1* mutation (31, 59).

NPM forms pentamers, that may assemble into decamers, through a conserved N-terminal domain (15, 33, 42, 51). This domain plays a crucial role in NPM interaction with many of its partners, e.g. with p14Arf or c-myc (11, 38, 45). NPM with C-terminal mutation has been reported to retain the ability to form oligomers (3, 37). Heterooligomer formation between NPMmut and NPMwt affects the localization of one another (7, 18). It results in a decrease of NPMwt concentration in the nucleolus and may thus cause a loss of function of the fraction of NPMwt delocalized into the cytoplasm (17). Interaction of NPMmut with tumor suppressors also leads to aberrant transport of these proteins into the cytoplasm and, presumably, to the loss or restriction of their biological function (13, 49). However, the role of the NPM mutation and of protein delocalization in AML initiation and in the treatment response has not been elucidated yet.

In general, two approaches are tested to prevent delocalization of NPMwt and of its interaction partners into the cytoplasm, resulting from their interaction with NPMmut. First, an inhibition of NPM nuclear exporter XPO1, and, the second, an impairment of the NPM oligomerization. The both approaches aim to reestablish the correct NPMwt localization (2, 62). Leptomycin B can block NPM transport into the cytoplasm, but it cannot be used for AML treatment owing to a high toxicity (19). Alternative second-generation XPO1 inhibitors, selinexor and eltanexor, are drugs with promising anticancer effect. Selinexor is currently being tested in a phase I clinical trial (25). Recently, its effect on complex formation between NPMmut and the transcription factor PU.1 with a key role in monocyte lineage differentiation have been demonstrated (30). In the present study, we focused on the second option, i.e. on the manipulation of NPM localization by an interference with NPM oligomerization.

The small molecule NSC348884 has been reported to prevent formation of NPM oligomers (61). It has been found to activate p53, to inhibit cell growth, and to trigger the apoptosis (2, 61). In pregnant mice, application of NSC348884 induced attenuation of p-Stat3 nuclear localization in the uterine luminal epithelium and decreased the number of implantation sites (36). Cell cycle arrest at G1 phase and viability suppression were observed after NSC348884 treatment of atypical teratoid/rhabdoid tumor (AT/RT) cell lines. Moreover, significant upregulation of *NPM1* gene, typical for AT/RT cells, was extensively reduced in the presence of NSC348884 (56). NSC348884 was also proved to reduce phagocytosis of myelin-debris by microglia. This reduction was ascribed to decreased K-Ras/Galectin-3 signaling possibly caused by NPM inhibition (64).

To study the effect of NPM oligomerization inhibitors, it was necessary to establish a reliable set of methods for detection of NPM oligomers in both cell lysates and live cells. Recently, we presented a method for monitoring NPM oligomerization in live cells based on a time-resolved fluorescence (FLIM-FRET) technique (35). In that work, point mutations in the NPM oligomerization domain, at Cysteine 21 (C21), have been introduced, as the mutation of C21 to aromatic hydrophobic residues had been reported to inhibit NPM oligomerization (37, 60). Surprisingly, we did not observe any effect of these mutations on NPM oligomerization in live cells (35). Although this result was also confirmed by immunoprecipitation, some changes induced by C21 mutations can be detected *in vitro* by native and semi-native electrophoresis, as we document in the present study. Enomoto *et al.* (16) determined specific residues and regions accountable for NPM oligomerization, for its nucleolar localization, and for p14Arf binding (16). These data disclosed that NPM variants lacking part of N-terminal domain were localized in the nucleoplasm and exhibited no ability to interact with other NPM molecules. As a test system in live cells, we thus created fluorescently labeled N-terminal NPM mutants with deletions of the first 25, 100 and 117 amino acids (Δ25, Δ100, Δ117), and we examined their oligomerization status by native electrophoresis, immunoprecipitation, and FLIM-FRET technique.

Having established a robust experimental system, we analyzed the effect of NSC348884 in both cell lysates and live cells. Effect of NSC348884 on NPM oligomerization, apoptosis, and cell adhesivity is presented, and putative mechanism of NSC348884 action is discussed.

## Material and Methods

### Cell culture and chemicals

Leukemia cell lines MV4-11, OCI-AML2, OCI-AML3, KG-1, and KASUMI-1 were purchased from DSMZ (Germany), HL-60 were from ECACC (GB), adherent cell lines HEK-293T and HeLa were kindly provided by dr. Š. Němečková (Department of Immunology, Institute of Hematology and Blood Transfusion) and dr. J. Malínský (Institute of Experimental Medicine, Czech Academy of Science), respectively. The cells were cultivated in growth media with fetal bovine serum (FBS), glutamine and antibiotics (all from Sigma-Aldrich) according to manufacturers’ recommendation: MV4-11, KG-1, HL-60, and HeLa in RPMI-1640/10% FBS, OCI-AML2 and OCI-AML3 in alpha-MEM/20% FBS, KASUMI-1 in RPMI-1640/20% FBS and HEK-293T in DMEM/10% FBS. Peripheral blood mononuclear cells (PBMC) originated from leukapheresis of hyperleukocytic AML patients. PBMC were separated using Histopaque 1077 (Sigma-Aldrich), washed with PBS and resuspended in RPMI-1640 with 10% FBS. The presence of C-terminal NPM mutation was detected by PCR and the mutation type was determined by sequencing (40) and confirmed by Western blotting and immunofluorescence using specific anti-NPMmut antibody (66). All patients signed informed consent to the use of their biological material for research purposes in agreement with the Declaration of Helsinki. The study has been approved by the Ethics Committee of the Institute of Hematology and Blood Transfusion of the Czech Republic. All cells were cultivated in 5% CO_2_ atmosphere at 37°C. Stock solution of 10mM NSC34884 was added to cell suspensions to final concentrations and times as indicated in the Results section.

### Plasmid construction and transfection

As we described in detail previously (6, 7), the gene for NPM was amplified from cDNA library (Jurkat cells, Origene) by PCR and inserted to vectors peGFP-C2 and pmRFP1-C2 (originally Clontech), designed for expression of protein chimeras with a fluorescent protein connected to the N-terminus of the target protein, by standard methods of molecular cloning. NPM mutants were constructed by PCR using extended primers targeting NPM1 sequence neighboring regions cut from the N-terminus or containing the mutated part of the exon 12 of the *NPM1* gene complemented with appropriate restriction sites (7). After amplification in E. coli, the plasmids with subcloned genes were purified with PureYield Plasmid Miniprep System (Promega) and transfected into adherent cell lines using jetPRIME transfection reagent (Polyplus Transfection).

### Cell lysis and western blotting

Cell lysis. As described previously (66), cells were washed with PBS and lysed depending on the intended application. For direct use in SDS-PAGE, the cells were lysed in Laemmli sample buffer (SB, 50mM Tris pH 6.8, 2% SDS, 100mM DTT, 10% glycerol), boiled at 95°C for 5min, centrifuged at 200.000g/4°C for 4h and the supernatant was stored at −20°C. For other applications, the cells were lysed in Lysis buffer (LB, 10mM Tris/Cl pH7.5, 150mM NaCl, 0.5mM EDTA, 0.5% NP-40, protease and phosphatase inhibitors) for 30min/4°C, centrifuged at 20.000g/4°C for 10min and supernatant was mixed 1:1 with the appropriate buffer.

Native and semi-native PAGE. Lysates were mixed with 2xnative buffer (NB, 50mM Tris pH6.8, 10mM DTT, 10% glycerol) and subjected without boiling to 7,5% AA Tris-glycin gel without SDS for native electrophoresis, or to the gel with SDS (2%) for semi-native electrophoresis.

Western blotting. Five to ten microliters of each sample were subjected to native or SDS-PAGE and transferred into PVDF or nitrocelulose membrane (BioRad). Mouse monoclonal antibodies against β-actin, GFP, dsRed, NCL, FBL, NPM (clone 3F291 for NPMwt+mut detection, clone E3 for NPMwt detection), and p14Arf were from Santa Cruz Biotechnology. All mouse primary antibodies were used at a dilution 1:100-1:500. Rabbit polyclonal antibody against NPMmut (pab50321, Covalab) was used at 1:2 000 dilution. Rabbit monoclonal anti-PAK1 (1:2 000, Abcam) and anti-PAK1-pSer144 (1:20 000, Abcam) and rabbit polyclonal antibodies and anti-Cofilin and anti-Cofilin-pSer3 (Cell Signalling Technology) were used for adhesion-related protein detection. Anti-mouse and anti-rabbit HRP-conjugated secondary antibodies were purchased from Thermo Scientific and used at concentrations 1:10.000-1:50.000. ECL Plus Western Blotting Detection System (GE Healthcare) was used for chemiluminescence visualization and evaluation by G-box iChemi XT4 digital imaging device (Syngene Europe). Alternatively, Alexa488-conjugated anti-rabbit and Alexa647-conjugated anti-mouse secondary antibodies (ThermoFisher) for simultaneous detection of NPMmut and NPMwt+mut were used.

### Immunoprecipitation

Immunoprecipitation using GFP- or RFP-Trap (Chromotek) was performed according to manufacturer’s instruction as described previously (7). Briefly, cells were harvested and washed with PBS, lysed in LB for 30min/4°C and centrifuged at 20.000g/4°C for 10min. The lysate was mixed with GFP/RFP-nanobeads and rotated for 1h/4°C. The beads were extensively washed with diluting buffer (10mM Tris/Cl pH7.5, 150mM NaCl, 0.5mM EDTA), resuspended in SB, boiled at 95°C for 10min and centrifuged 20.000g/4°C for 10min. Supernatant was stored at −20°C until used for SDS-PAGE.

### Live-cell imaging

The cells were seeded in the 2,5mm culture dish with glass bottom (Cellvis) for 24h and then transfected with plasmids containing fluorescent variants of the desired genes. After another 24h, the transfected cells were observed under confocal laser scanning microscope FluoView FV1000 (Olympus Corporation) using 543nm excitation for RFP fluorescence and 488nm excitation for GFP and for differential interference contrast (DIC) observation. UPlanSAPO 60x NA1.35 oil-immersion objective was used for imaging. For long-term monitoring, the culture dish was sealed by parafilm to prevent CO_2_ leakage and it was placed into microscopy chamber tempered to 37°C. NSC348884 was added just before the start of the measurement. Fluorescence images were processed by the FluoView software FV10-ASW 3.1.

### Lifetime imaging and acceptor bleaching

The apparatus used for lifetime imaging is described in detail elsewhere (32). Briefly, we used inverted IX83 microscope equipped with a FV1200 confocal scanner (Olympus, Germany), cell cultivation chamber (Okolab) and FLIM add-ons from PicoQuant. Fluorescence was excited by a pulsed diode laser (LDH-DC-485, 485 nm, PicoQuant) running at 20 MHz repetition rate. Light was coupled to the microscope by a single-mode optical fiber and reflected to the sample by 488 nm long-pass dichroic mirror (Olympus). Typically, UPLSAPO 60XW NA 1.2 water-immersion objective (Olympus) was used for imaging. Fluorescence was directed via multimode optical fiber to a cooled GaAsP hybrid PMT (PicoQuant) through the 520/35 bandpass filter (Semrock). Signal was processed by the TimeHarp 260-PICO TCSPC card and the SymPhoTime64 software (both PicoQuant). To avoid pile-up artifacts, the data collection rate at brightest pixels was kept below 5% of the excitation frequency. FLIM images were collected in a few minutes with the excitation power around 0.1 μW. Acceptor photobleaching was done by a 561 nm semiconductor CW laser with a multi-mW power at the focal point. All experiments were done at 37°C.

### Lifetime Data processing

Lifetime images were generated in the SymPhoTime64 by the “fast-FLIM” method when pixel lifetimes were calculated by a method of moments (55). Specifically, pixel lifetimes *τ*_*avg*_ were determined as the difference between the barycenter of the fluorescence decay and the time-offset *t*_*offset*_ of the steepest growth of the decay at each pixel:

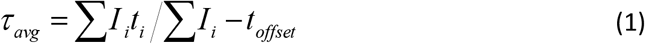

where *I*_*i*_ is a decay intensity at time *t*_*i*_. Exported FLIM images were further processed and visualized by the Fiji software (68). An accurate analysis of the cumulative decays from larger area of interest was done by the least-squares reconvolution also in the SymPhoTime64. Fluorescence was assumed to decay multiexponentially according to the formula:

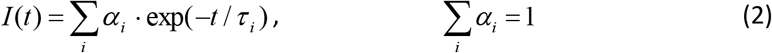

where *τ*_*i*_ and *α*_*i*_ are the fluorescence lifetimes and the corresponding amplitudes, respectively. Typically, two decay components were sufficient for acceptable fit. The intensity-weighted mean fluorescence lifetime was calculated as:

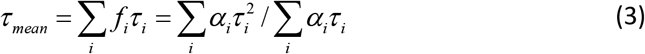

where *f*_*i*_ are fractional intensities of the *i-*th lifetime component:

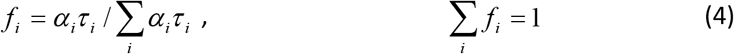

### Electrical cell-substrate impedance sensing (ECIS)

Impedance measurements were performed using the ECIS Zθ device (Applied Biophysics). The wells of 8W10E+ plates were filled with 200 µl culture medium and the baseline was monitored for several hours before cell addition. HeLa or 293T cells were seeded at 120.000 cells/well and monitored overnight, the inhibitors were added after 20 to 24 h. One well from each plate was left empty (medium only), and the signal from this well was used as the baseline for the other wells of the same plate. The instrument automatically decomposes the impedance signal into resistance and capacitance. As the course of capacitance at 64 kHz mirrored that of resistance at 2 kHz, the observed evolution of the resistance signal reflects changes in the cell-surface contact area. The ECIS records were exported to Microsoft Excel and processed using the GraphPad Prism software: the background was set to zero at a time point shortly before cell seeding, and the baseline (empty well) was subtracted. The curves shown in the graphs represent the averages from replicate wells, which were run in parallel.

### Statistical analyses

As described in our previous work (7), the majority of experiments were performed using cell lines and repeated until the observed differences between groups reached statistical significance. A p-value of 0.05 or lower was pre-set to be indicative of a statistically significant difference between groups compared. In diagrams, arithmetic means of replicates of all experiments were plotted with SD error bars. Significance levels (p values of ANOVA or Student’s t-test) were determined using InStat Software (GraphPad Software).

## Results

### Interaction and stability of NPM with C21 point mutation

NPM oligomerization is mediated by the N-terminal NPM part whereas the mutation, which is associated with acute myeloid leukemia (AML) and which drives NPMmut into the cytoplasm, occurs at the C-terminus. This mutation does not abrogate the NPM oligomerization ability (3). Partial delocalization of NPMwt into the cytoplasm is caused by heterooligomer formation in cells expressing NPMmut (3, 7). The native electrophoresis performed under reducing conditions (termed the semi-native electrophoresis, see Material and Methods) revealed that the stability of NPM oligomers was slightly lowered in the C-terminal mutants (66). Therefore, therapy based on the disruption of NPM oligomerization might be beneficial for AML patients with *NPM1* mutation. A reliable method for oligomerization monitoring is required in screens for potential drugs affecting the oligomerization. To establish a positive control and to validate the method, we tested mutations reported to inhibit NPM oligomerization.

Point mutation in C21 was shown to be important for NPM oligomerization (60). However, we have recently demonstrated that in live cells, oligomeric state of NPM is not affected by this mutation (35). In addition, immunoprecipitation revealed the interaction of the endogenous NPM with eGFP-labeled NPM with point mutation at C21 (G_C21). C21 was substituted either to Ala (G_C21A), or to Phe (G_C21F). In the present work, we used fluorescence microscopy and native electrophoresis to characterize in detail the impact of these C21 mutations. Both G_C21 mutants exhibited nucleolar localization, identical to that of NPMwt (Fig. 1). Furthermore, the mRFP1-labeled NPM_C21 (R_C21) was found in the cytoplasm of HEK-293T (293T) cells co-transfected with eGFP-labeled NPMmut (G_NPMmut). The fraction of cells exhibiting mRFP1 signal in the cytoplasm was comparable for all R_NPM variants (Fig. 1). Both C21 mutants therefore seem to form heterooligomers with NPMmut alike NPMwt. To follow the findings on the C-terminal mutants, we examined native lysates from cells expressing G_C21A or G_C21F in acrylamide gels with and without SDS (Fig. 2). Whereas C21A exhibited a high-MW band (presumably oligomers) identical to that of NPMwt under native conditions, the band from C21F was located at the position corresponding to the weak lower-MW fraction of NPMwt or NPMmut (presumably monomers). In these experiments, the band from endogenous NPM oligomers served as a loading and position control. The results from semi-native electrophoresis in Fig. 2 show markedly increased monomer/oligomer ratio of C21A compared to NPMwt and absence of C21F oligomers. These results suggest that although NPM oligomerization seems to be unaffected by the C21 point mutation in live cells, the stability of the oligomers is considerably attenuated. Similar results were obtained with mRFP1-labeled variants. Interestingly, mRFP1-labeled proteins displayed slightly lower mobility in the native conditions. Moreover, R_NPM oligomers were found to be somewhat more stable compared to G_NPM ones (Fig. 2b).

**Fig. 1:**
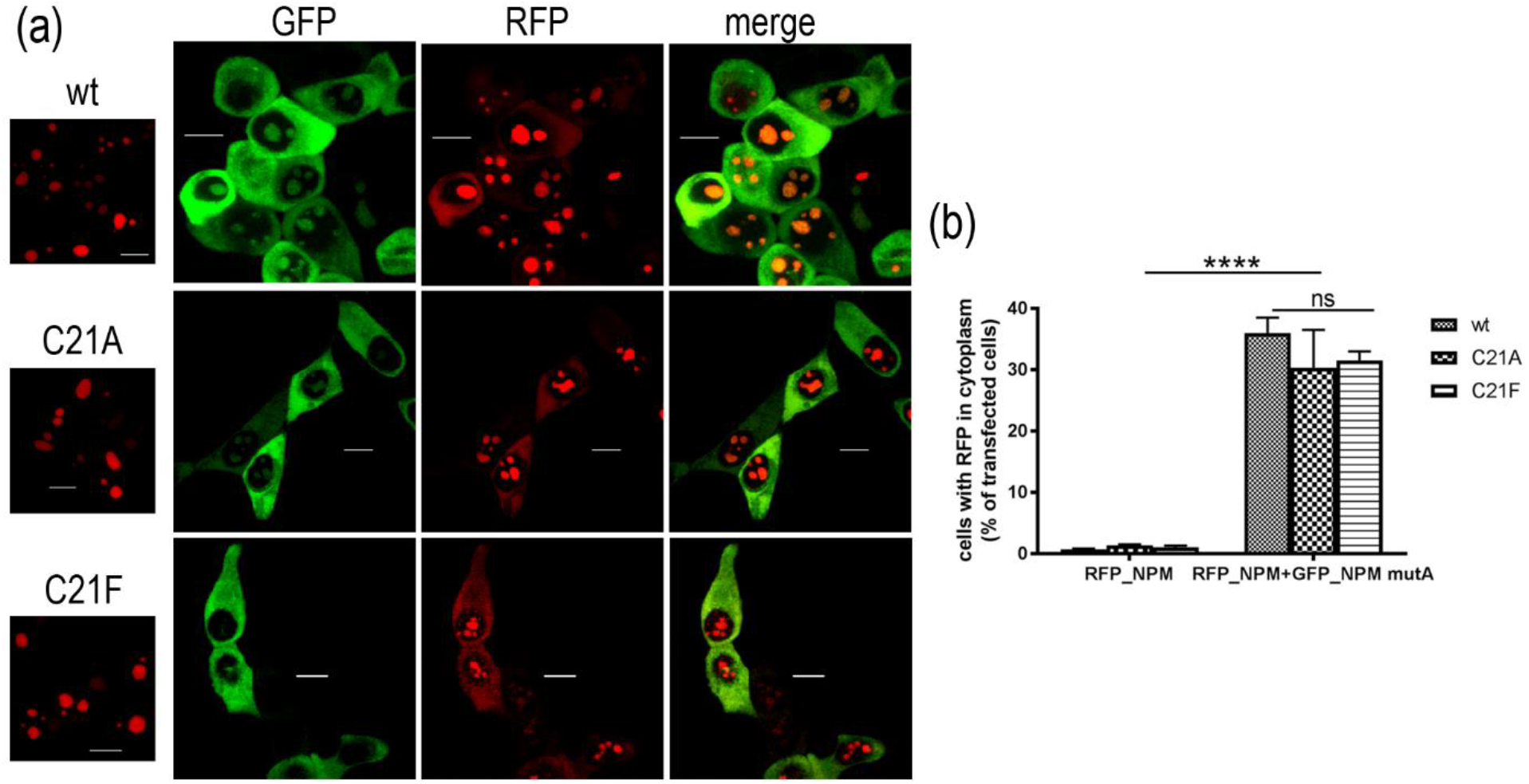
Interaction between NPMmut and C21 point mutants. (a) Left: localization of R_NPMwt/C21A/C21F in single transfected 293T cells. Right: 293T cells co-transfected with G_NPMmut (green) and R_NPM variants (red). Red signal in the cytoplasm and green signal in nucleoli witness for an interaction between NPMmut and NPMwt/C21A/C21F. (b) Statistical evaluation of subcellular localization of NPMwt and C21. Fraction of transfected 293T cells displaying red signal from the cytoplasm in single transfected cells (left) and in cells co-transfected with G_NPMmut (right). Error bars represent ±SD of at least 3 independent experiments, p<0,0001 (****) for the differences between the co-transfected and the single-transfected cells.

**Fig. 2:**
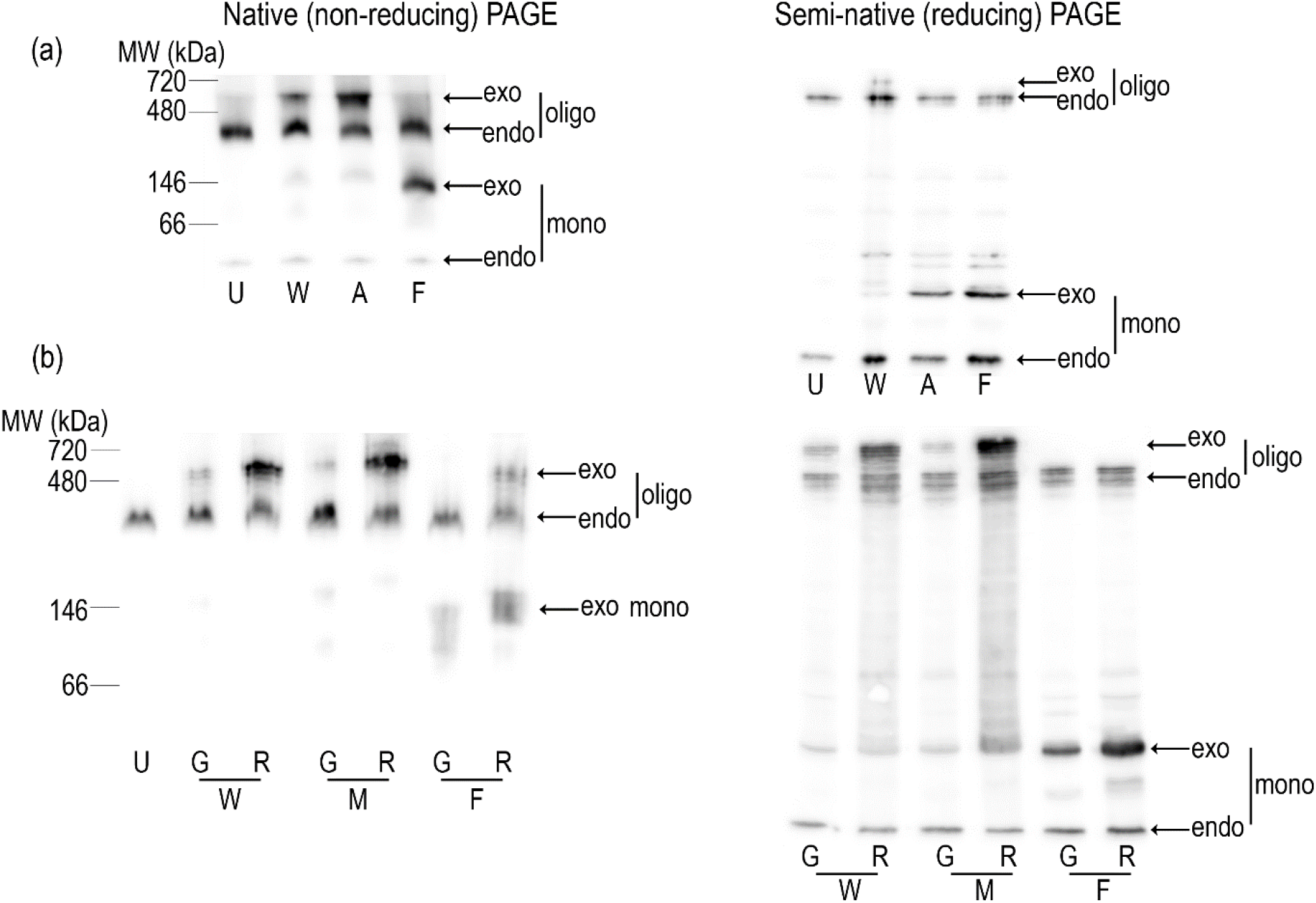
Effect of C21 mutations on the stability of NPM oligomers. Native (left) and semi-native (right) PAGE analysis of lysates from untransfected (U) 293T cells or cells transfected with NPMwt (W), C21A (A), C21F (F) or NPMmut (M). (a) Stability of oligomers formed by various G_C21 constructs is affected by the reducing PAGE conditions. (b) Position of bands and oligomer stability depend on the fluorescent label used (G or R: eGFP- or mRFP1-labeled proteins). The figures show representative examples from repeated experiments.

We have not observed any changes in expression and in oligomerization state of the endogenous NPMwt in cells containing C21 mutants. Therefore, we further investigated the effect of C21F substitution on the stability of exogenous heterooligomers formed by a mixture of fluorescently labeled C21F and NPMwt. 293T cells were alternatively transfected with single fluorescent variants of C21F and NPMwt, or co-transfected with the combination of both. As shown in Fig. 3 and Fig. S1, the exogenous NPMwt monomer/oligomer ratio is strongly affected by the presence of C21F. In the native blots, band intensities attributed to the C21F oligomers and to the NPMwt monomers are both considerably higher in traits corresponding to co-transfected sample than the values in traits from single transfected cells. Occurrence of NPM molecules either in oligomer (high-MW) or monomer (low-MW) bands likely depends on the NPMwt/C21F participation in the heterooligomers. Interestingly, endogenous NPM monomer/oligomer ratio seems to be unaffected by the presence of any exogenous NPM form. Altogether, despite the C21F mutation does not abrogate NPM oligomerization in live cells, it evidently attenuates the oligomer stability.

**Fig. 3:**
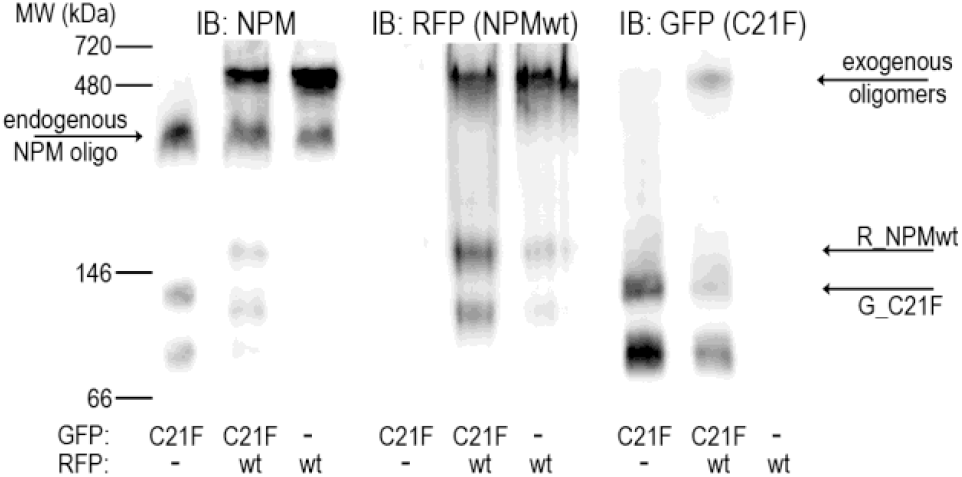
Formation of heterooligomers containing NPM with C21F substitution and NPMwt. Western blots of native PAGE of samples from 293T cells transfected with R_NPMwt (wt), G_C21F (C21F), and with their combination. Similar results were obtained with the inverse tagging combination, i.e. with G_NPMwt and R_C21F.

### Localization and oligomerization properties of NPM N-terminal deletion mutants

In view of the fact, that the C21-point mutations cause no detectable changes of the NPM oligomerization in live cells, we searched for other modifications of the NPM oligomerization domain in order to validate detection methods for the oligomerization disruption. Numerous NPM N-terminal deletion mutants were reported to lose the oligomerization ability and the nucleolar localization depending on the extent and specificity of the deleted region (16). To determine the N-terminal region required for NPM oligomerization, we constructed three N-terminal mutants having the first 25 (Δ25), 100 (Δ100), or 117 (Δ117) amino acids deleted. Then we analyzed their subcellular localization and oligomerization characteristics. Our confocal imaging experiments have revealed that all the truncated NPM forms reside both in the nucleoli and in the nucleoplasm (Fig. 4). As expected, the largest deletion resulted in an increased accumulation of the mutant in the nucleoplasm.

**Fig. 4:**
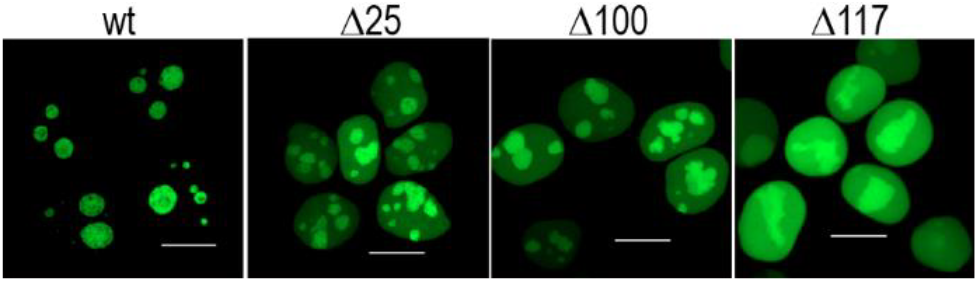
Significance of the N-terminus for NPM localization. Diminished nucleolar accumulation and increased amount in the nucleoplasm of Δ25, Δ100 and Δ117 compared to WT.

To monitor NPM complex formation, lysates from 293T cells transfected with fluorescently labeled deletion mutants were subjected to electrophoresis. As expected, inability of the truncated proteins to form oligomers was reflected by an absence of the high-MW bands under native and semi-native conditions (Fig. 5a). Further we investigated the stability of mixed NPM oligomers containing exogenous NPMwt and selected deletion mutants. 293T cells were co-transfected with plasmids ensuring expression of Δ117 and NPMwt and the lysates were analyzed (Fig. 5b and S2). Bands from oligomeric NPMwt complexes are unaffected by the presence of Δ117 in the native as well as in the semi-native immunoblots. The result suggests that, in contrast to the C21 mutants, the presence of the deletion mutants did not affect the stability of the NPMwt oligomers. Again, expression of the endogenous NPMwt remained unchanged. To analyze the interaction potential of the N-terminal deletion mutants, 293T cells were transfected with plasmids encoding for GFP-labeled NPMwt, Δ25, Δ100 or Δ117. Then co-immunoprecipitation using GFP-Trap was performed. Surprisingly, interaction of the endogenous NPM with any deletion mutant (i.e. except the plasmid encoding for free eGFP) was found although the amount of co-precipitated endo-NPM was slightly lower compared to the G_NPMwt precipitates (Fig. 5c). Simultaneously, level of co-precipitated nucleolin (NCL), which interacts with the NPM region near its C-terminus (44) and does not interact with the AML-related NPMmut (66), was higher in precipitates of the truncated NPM forms. Interestingly, the level of another co-precipitated nucleolar protein, fibrillarin (FBL), also positively correlated with the extent of N-terminus deletion. Finally, the tumor suppressor p14Arf, which is known to interact with the N-terminal NPM domain (16), clearly co-precipitated only with the G_NPMwt.

**Fig. 5:**
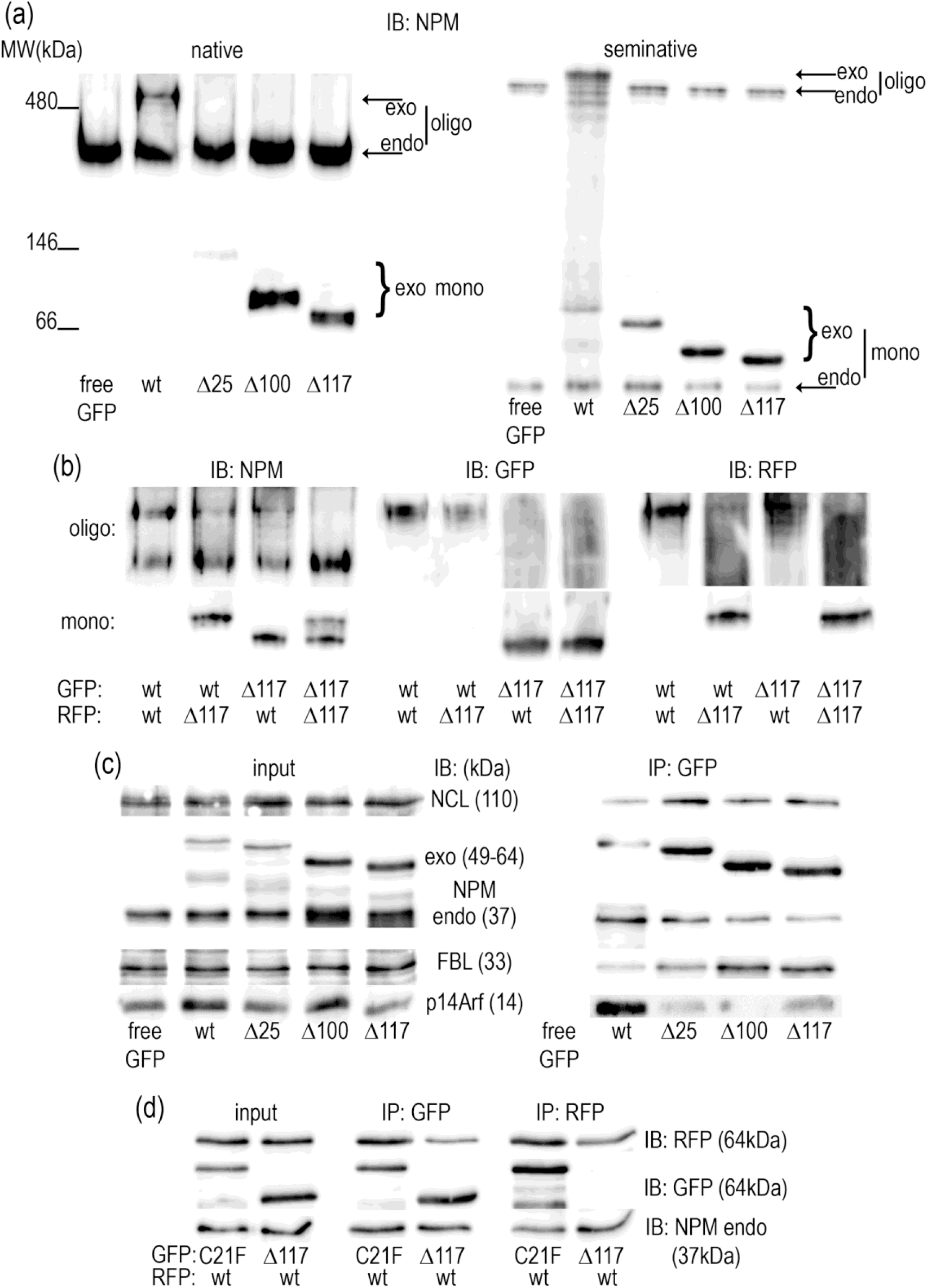
Significance of the N-terminus for NPM oligomerization. (a) NPM expression in native and semi-native PAGE of 293T cells transfected with free eGFP and eGFP-tagged truncated NPM variants. (b) Native PAGE of 293T cells co-transfected with combinations of Δ117 and NPMwt illustrating absence of interaction between Δ117 and NPMwt. (c) Interaction of truncated NPM forms with endogenous proteins. Lysates from 293T cells expressing GFP-labeled NPMwt, Δ25, Δ100 and Δ117 were subjected to immunoprecipitation and the levels of co-precipitated interaction partners were investigated. (d) eGFP/mRFP1-immunoprecipitation from 293T cells co-transfected with R_NPMwt and G_C21F or G_Δ117: asymmetric results of precipitation from the sample containing the truncated NPM form.

To evaluate the utility of the method for detection of the complexes containing both NPMwt and the deletion mutants, we performed GFP/RFP-immunoprecipitation from the cells co-transfected with R_NPMwt and G_Δ117. The co-transfection of R_NPMwt and G_C21F served as the interacting control. Whereas NPMwt was clearly detected in samples obtained by precipitation of the deletion mutant, the vice versa co-precipitation failed (Fig. 5d). Identical results were obtained for combination NPMwt+Δ100. No interaction was found between two color variants of the N-terminal mutant (Fig. S3).

The Förster resonance energy transfer (FRET) is a robust spectroscopic method for evaluation of the donor-acceptor proximity in protein complexes. Increased FRET upon the complex formation is reflected in the decrease in the donor fluorescence lifetime. We used fluorescence lifetime imaging (FLIM) to evaluate its capability to detect complexes between fluorescently tagged truncated proteins and NPMwt in live cells. Specifically, the FRET between eGFP donor and mRFP1 acceptor attached to NPMwt and Δ117 (or Δ100) was examined. Presence of FRET within the complex was detected by the acceptor photobleaching: an increase in the donor lifetime upon the acceptor photodestruction is a strong positive indicator of the mixed-complex formation. Results are shown in Fig. 6. Panels A and B show the initial localization of the green and red signal within the cells, panels C and D the intensity ratio I_red_/I_green_ before and after the photobleaching, respectively. Corresponding FLIM images and lifetime histograms from the analyzed nucleolar area are shown in the panels E, F and G, respectively. As expected, a significant lifetime increase resulting from the FRET cancellation was detected after photobleaching in cells co-transfected with G_NPMwt and R_NPMwt. On the contrary, virtually no effect was observed in cells transfected with the G_Δ117+R_Δ117 mutants. In accord with the literature (16) and with our precipitation data, this result indicates inability of the Δ117 mutants to interact with each other and to form multimers. This protein pair can therefore serve as a negative control for the multimer formation. As seen from Fig. 6, in samples containing combination of the Δ117 deletion mutants with NPMwt, the GFP-lifetime sligthly, but still visibly, increases upon the acceptor photobleaching. The change corresponds with the results of our electrophoretic experiments and suggests some amount of the mixed multimer to be present in the cell. The statistical analysis from multiple experiments (n=3-5) is presented in Fig. 6H, where cells transfected with G_Δ117 (the donor only transfection) served as a negative control. The figure clearly proves oligomerization of NPMwt. Oligomerization of Δ117 deletion mutants is clearly undetected and the presence of mixed NPMwt+Δ117 multimers is at the significance limit. Interestingly, the statistical evaluation also suggests the asymmetric character of interaction between NPMwt and Δ variants. These results are in a nice agreement with the immunoprecipitation.

**Fig. 6:**
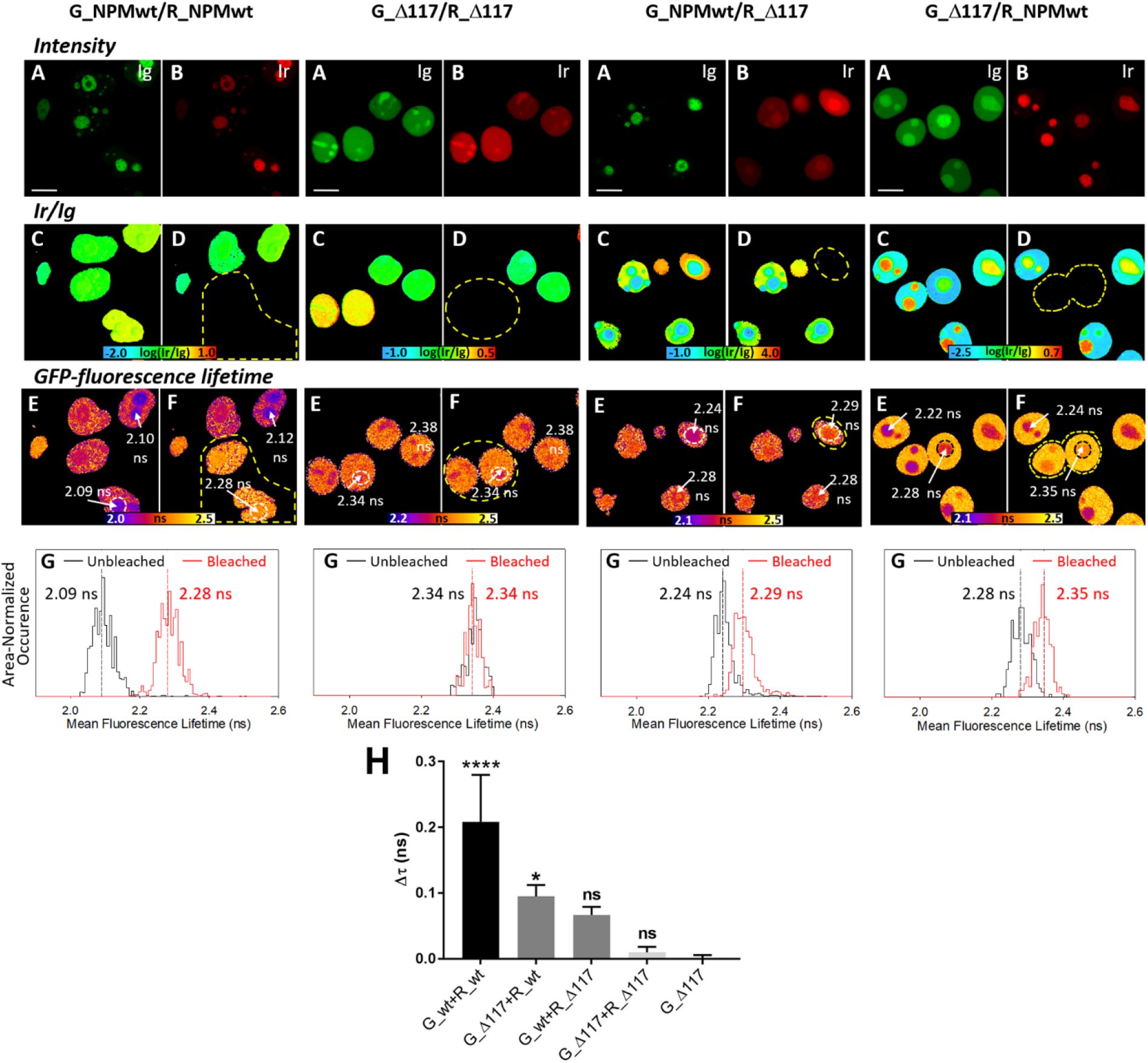
Effects of deletion in the N-terminal domain of NPM on its oligomerization in living 293T cells studied by FLIM-FRET. Interaction of eGFP- and mRFP1-labeled NPMwt causes shortening of the eGFP-fluorescence lifetime (τ) in co-transfected cells. After mRFP1-photobleaching, τ becomes prolonged, which confirms the G_NPMwt/R_NPMwt interaction. No lifetime change after mRFP1 photobleaching suggests absence of interaction between G_Δ117 and R_Δ117. A, B: Initial localization of the green and red signal within the cells; C, D: The intensity ratio of I_red_/I_green_ before and after the photobleaching, respectively. E, F: Fluorescence lifetime distribution of eGFP before and after the mRFP1-photobleaching; G: Lifetime histograms from the analyzed nucleolar area; (H): One-way ANOVA analysis of the eGFP fluorescence lifetime change after acceptor photobleaching. Changes are compared to the negative control G_Δ117. Error bars represent ±SD of at least 3 independent experiments (****, p<0.0001; *, p<0.05).

### Effect of NSC348884 on cell viability, apoptosis and NPM oligomerization

The small molecule NSC348884 was reported to interfere with NPM oligomerization in solid tumor cell lines (61) as well as in leukemia cells (2). Furthermore, it was proved to inhibit proliferation, to upregulate p53 and to trigger apoptosis (2, 61). We thus investigated the influence of NSC348884 treatment in a panel of leukemia cell lines complemented with HeLa and 293T cells.

Prior to characterization of the NSC348884 effect on NPM oligomerization, we performed the analysis of cell viability and apoptotic markers. The cell viability in the presence of NSC348884 was monitored by propidium iodide (PI) exclusion test (Fig. 7a). Caspase-3 fragmentation as well as changes of p53 expression were investigated by immunoblotting to assess the extent of apoptosis in NSC348884-treated cells (Fig. 7b). For the majority of the cell lines, the EC50 value were within the interval from 2 to 10 μM. The viability drop correlated with increased caspase-3 fragmentation indicating the onset of apoptosis (Fig.7b). Simultaneously, NSC348884-induced increase in the p53 level was detected in some of the cell lines possessing wild-type p53. Contrarily to previously reported results (2), the majority of cell lines with NPMwt was more sensitive than the cell line with NPMmut (OCI-AML3). Comparable sensitivity (from caspase-3 fragmentation) to NSC348884 treatment was found also for the primary cells of AML patients regardless of their NPM mutational status (Fig. 7c). Unexpectedly, native PAGE experiments revealed no influence of NSC348884 on NPM oligomerization (Fig.8). Endogenous NPM oligomers were found to be stable in KG-1, HL-60, MV4-11, and HeLa cell lines, which exhibited extensive apoptosis after NSC348884 treatment, as well as in OCI-AML2, OCI-AML3 and 293T, which were substantially more resistant to the treatment.

**Fig. 7:**
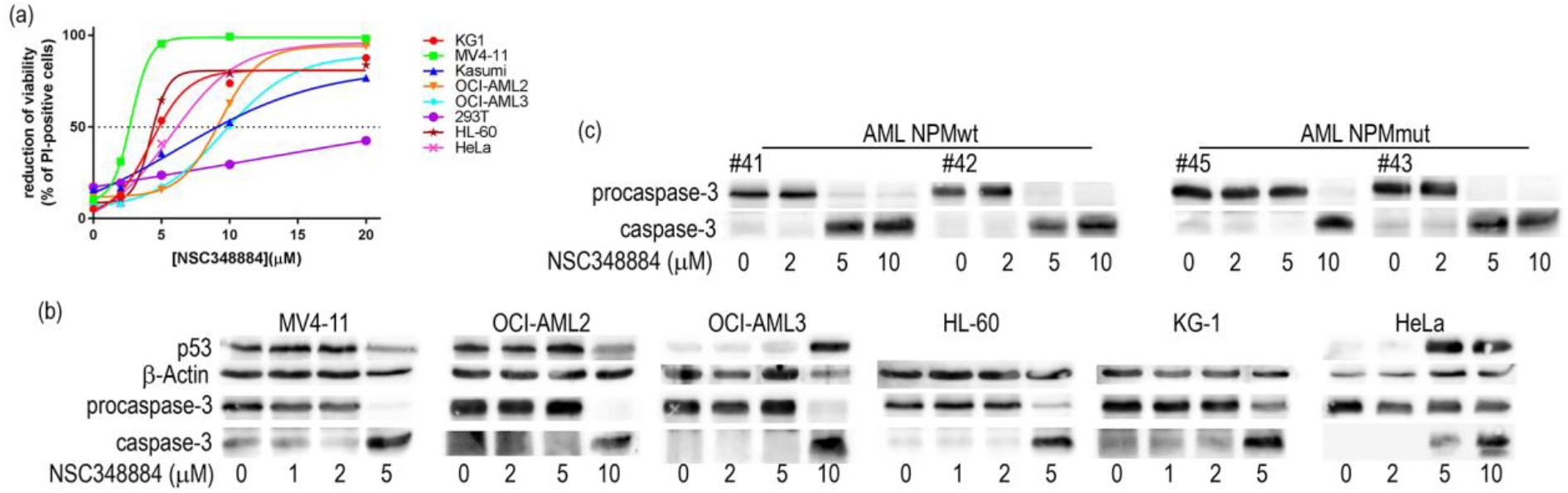
The effect of 24h NSC348884-treatment on cell viability and apoptosis. (a) Cell viability monitored by propidium iodide exclusion: each point represents the mean value of 3-10 independent experiments. (b-c) Representative blots of caspase-3 fragmentation and p53 expression in cell lines (b) and in primary AML cells (c). β-Actin levels serve as a loading control.

To further investigate effect of NSC348884 *in vivo*, we co-transfected 293T and HeLa cells with R_NPMwt and G_NPMmut. Then the cytoplasmic localization of R_NPMwt was monitored for 2 hours after addition of 10μM NSC348884 (Fig. 9). In agreement with our previous results (6, 7), detectable fraction of R_NPMwt was found in the cytoplasm of both cell lines at the starting time point. Lower cytoplasmic fraction of R_NPMwt in HeLa cells (compared to 293T) likely results from a higher endogenous NPM level (7, 47). Importantly, the cytoplasmic localization of R_NPMwt remained unchanged for at least 2h after the treatment, suggesting independence of NPM oligomerization on the presence of NSC348884 *in vivo* (Fig. 9). Simultaneously, there was an obvious effect of NSC348884 on cell-surface adhesivity. The effect is clearly visible in transmitted light images (DIC). Whereas the 293T cells progressively rounded and finally lost their contact with the glass surface of the culture dish, the HeLa cells detached from the surface individually. In any case, mitotic cells that detached from the surface for cell division have never re-adhered.

**Fig. 8:**
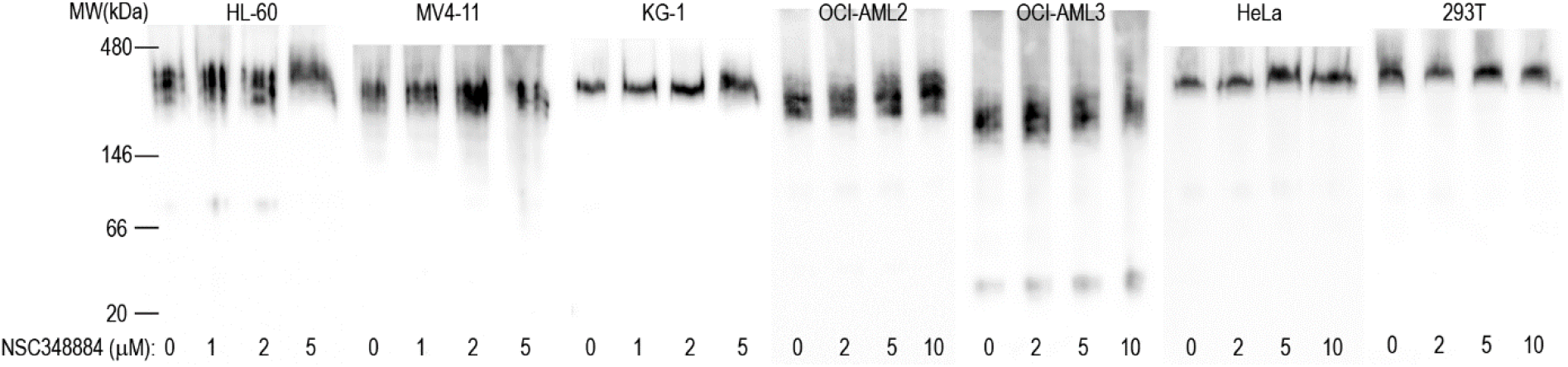
Effect of NSC348884 treatment on NPM oligomerization in leukemia cells. Native PAGE, representative blots: effect of 24h NSC348884 treatment on NPM oligomerization in various leukemic cell lines as well as in adherent HeLa and 293T cells.

**Fig. 9:**
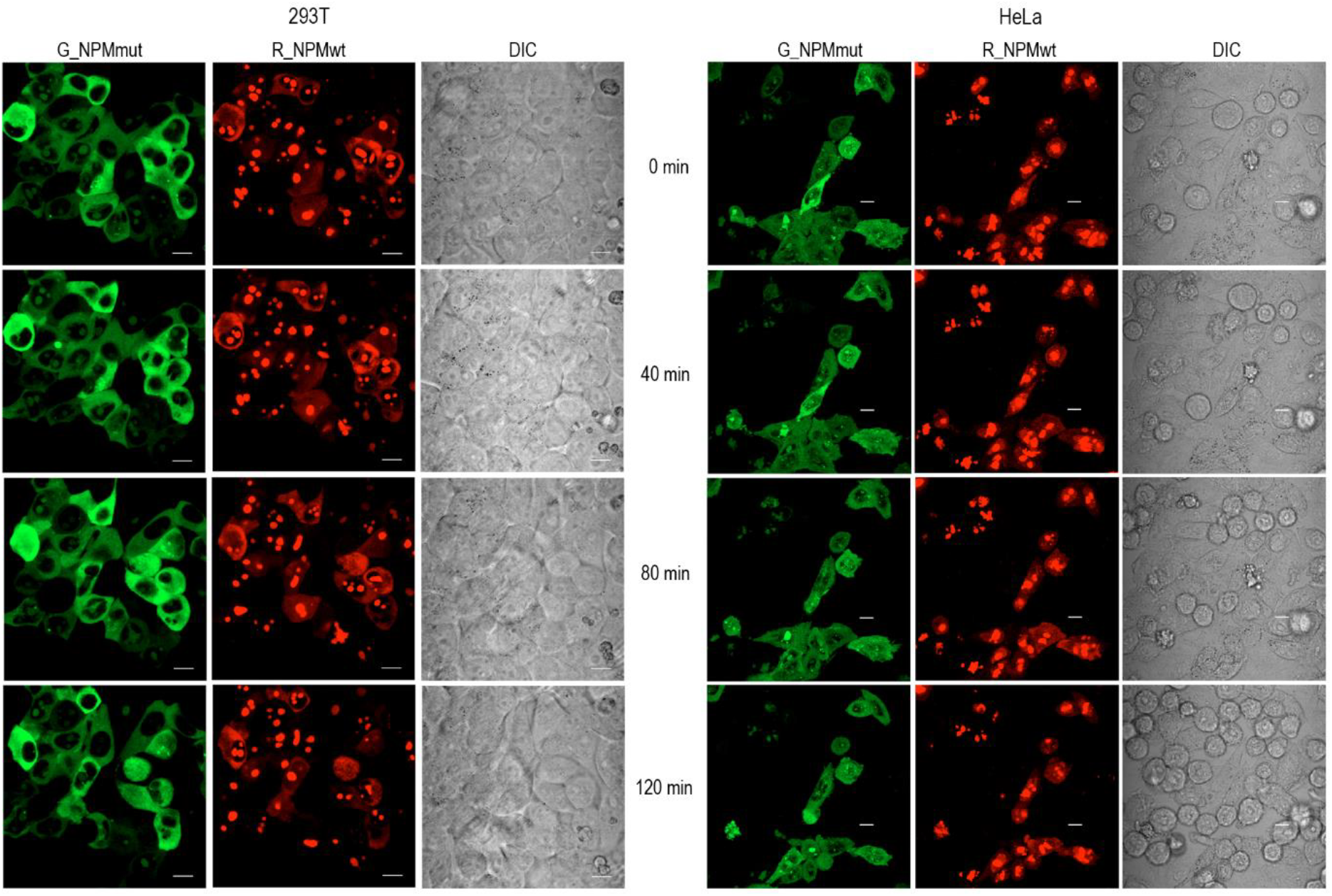
Time course of NSC348884-induced effects on the cell morphology. Effect of 10 μM NSC348884 was monitored in cells co-transfected with R_NPMwt (red) and G_NPMmut (green). The presence of the red signal in the cytoplasm and the cell morphology (DIC) were analyzed under confocal microscope. Left: 293T cells; right: HeLa cells. Scale bar represents 10μm.

The oligomerization of fluorescently labeled NPM was finally tested by the native PAGE and by immunoprecipitation in cell lysates and by FRET in live cells. First, we tested whether the low-MW band attributed to NPM monomers appears in native lysates of 293T cells expressing a combination of G_NPMmut and R_NPMwt after the NSC348884 treatment. Lysate from cells co-expressing G_NPMmut and weakly oligomerizing R_C21F was used to mark the position of the low-MW band (Fig. 10a and S4). No difference between the control and the NSC348884-treated sample was found either under native or semi-native conditions. Similar results were obtained from cells co-transfected with alternative combinations, i.e. with G_NPMwt+R_NPMwt or with G_NPMmut+R_NPMmut (Fig. 10b). Identical samples were afterwards subjected to immunoprecipitation (GFP- and RFP-Trap). All the exogenous NPM forms as well as the endogenous NPM were detected in all GFP- and RFP-precipitates regardless the NSC348884 addition (Fig. 11). In agreement with our previous work (66), control experiment revealed that NCL co-precipitated with NPMwt and it did not co-precipitate with any form of NPMmut. Again, the NPM-NCL interaction was not affected by the presence of NSC348884 in any experiment.

**Fig. 10:**
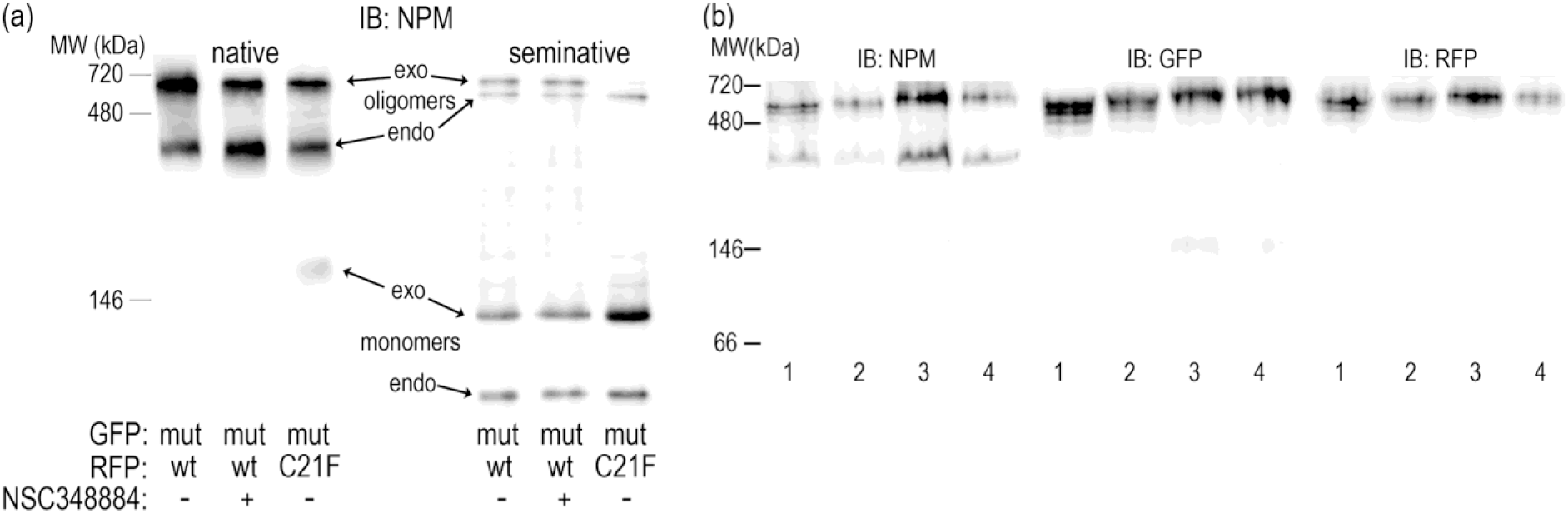
Native and semi-native PAGE from 293T cells transfected with various fluorescent variants of NPM and treated with 10 μM NSC348884 for 24h. (a) Cells co-transfected with G_NPMmut and R_NPMwt (lanes 1 and 2) or R_C21F (lane 3). Endogenous NPM, R_NPMwt and G_NPMmut oligomers detected in untreated (lanes 1 and 3) and NSC348884-treated (lane 2) cells. (b) Native PAGE, G_NPMwt+R_NPMwt (1, 2) and G_NPMmut+R_NPMmut (3, 4) in control (1, 3) and NSC348884-treated (2, 4) cells.

**Fig. 11:**
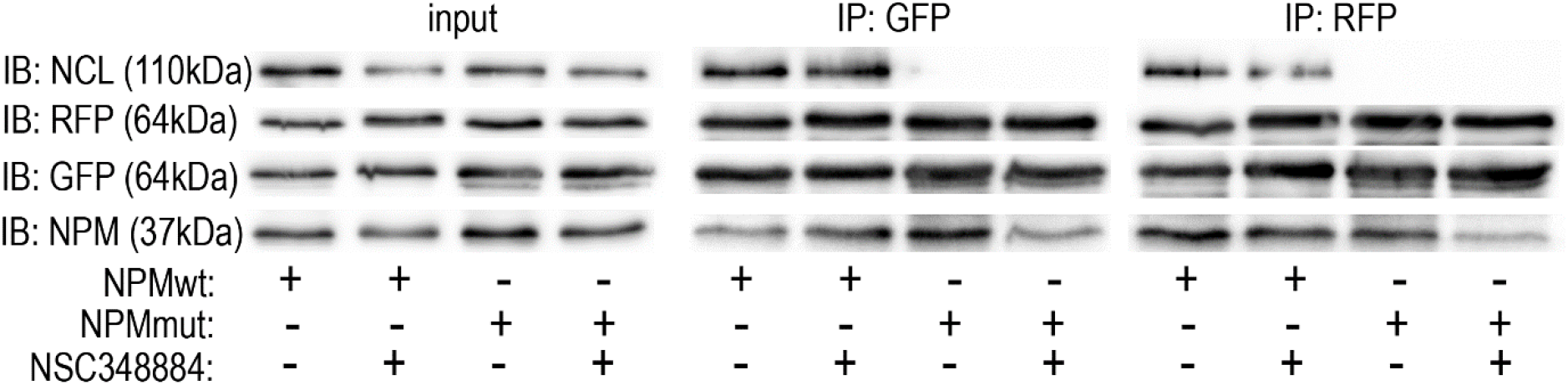
Interaction potential of NPM in NSC348884-treated cells. Immunoprecipitation from 293T cells co-transfected with G_NPMwt+R_NPMwt (NPMwt) or G_NPMmut+R_NPMmut (NPMmut) in control and NSC348884-treated sample (10μM NSC348884 for 24h). All exogenous forms as well as the endogenous NPM were detected in all eGFP- and mRFP1-precipitates regardless the NSC348884 addition. Nucleolin (NCL) was detected in NPMwt-precipitates only, independently of the NSC348884 treatment.

Oligomerization in live cells was independently tested by the resonance energy transfer. As seen from Fig. 12, FLIM-FRET experiments reveal unchanged eGFP fluorescence lifetime upon NSC348884 treatment of cells co-transfected with donor- and acceptor-labeled NPMwt and NPMmut. Prolonged eGFP (donor)-fluorescence lifetime after mRFP1 (acceptor)-photobleaching confirmed the complex formation in control cells without NSC348884 (column 1 and 2). The NSC348884-treatment did not affect the lifetime pattern (column 3) and the second round of the acceptor bleaching confirmed persistence of heterooligomers despite the presence of NSC348884 (column 4). Lower FRET extent in the NPMmut co-transfected cells (the second row) is likely a result of lower cytoplasmic NPMmut concentrations. Nevertheless, presence of FRET is still detected. NSC348884 activity resulting in cell rounding and loss of their contact with the glass surface is visibly documented by the morphology screening during the FLIM experiments (columns 5, 6), similarly as in Fig. 9. No lifetime change following mRFP1-photobleaching or NSC348884 treatment was detected in the control sample (Fig. S5), i.e. in cells expressing two color variants of NCL, where FRET was not detected (35). Altogether, FLIM-FRET experiments confirmed that NSC348884 does not affect NPM oligomerization, although it influences apoptosis and cell adhesion.

**Fig. 12:**
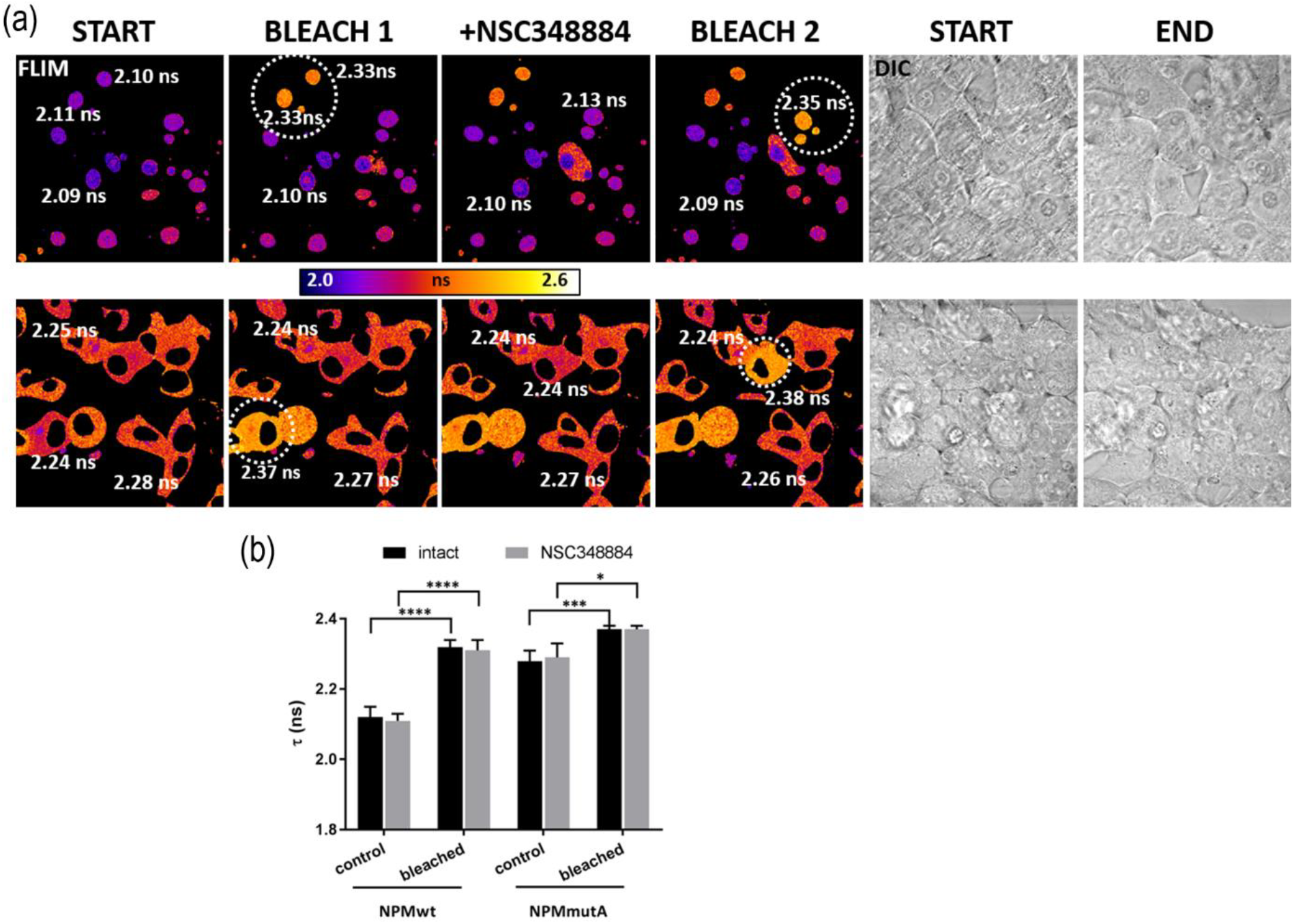
NPM oligomerization after NSC348884 treatment in live cells. (a) FLIM-FRET analysis of the eGFP-fluorescence lifetime (τ) after 2h of 10μM NSC348884 action on cells co-transfected with red and green variants of NPMwt (upper row) or NPMmut (lower row). White numbers: Fluorescence lifetime measured in individual cells. Dashed circles: region of mRFP1-photobleaching. Simultaneous cell morphology screening by DIC documents cell rounding induced by NSC348884. (b) Statistical evaluation of τ values before (control) and after (bleached) the mRFP1 photobleaching in intact (black bar) and NSC348884-treated (grey bar) cells. Student’s t-test of “control” vs. “bleached” values: ****p<0,0001, ***p<0,001,*p<0,05.

To further investigate the mechanism of NSC348884 action, we monitored the observed adhesivity changes more thoroughly with help of Electrical Cell-Substrate Impedance Sensing (ECIS) technique. As ECIS allows for real-time monitoring of cell contact with the surface of the sample well, we were able to follow the time course of the adhesivity decrease after NSC348884 addition (Fig 13). We have reported previously, that inhibition of Src family kinases by dasatinib resulted in a rapid drop of ECIS signal, which corresponded to cell shrinkage, and we thus used dasatinib as a reference compound (65). In both adherent cell lines, 293T and HeLa, NSC348884 induced large, dose-dependent changes in the resistance signal, which were similar to those produced by IPA-3 (27), an inhibitor of p21-activated kinases (PAK). PAK are key regulators of adhesion signaling, which have been proposed as therapeutic targets in different kinds of cancer including leukemias (10, 23). We thus analyzed possible effect of NSC348884 on expression and activity of PAK1, as well as of Cofilin, which governs actin remodeling during changes of cell shape. Indeed, Ser144 phosphorylation of PAK1 reporting on its kinase activity was reduced after 2h of NSC348884 treatment whereas total PAK1 expression remained unchanged (Fig. 14). Simultaneously, inactivating phosphorylation at Ser3 of Cofilin was detected.

**Fig.13:**
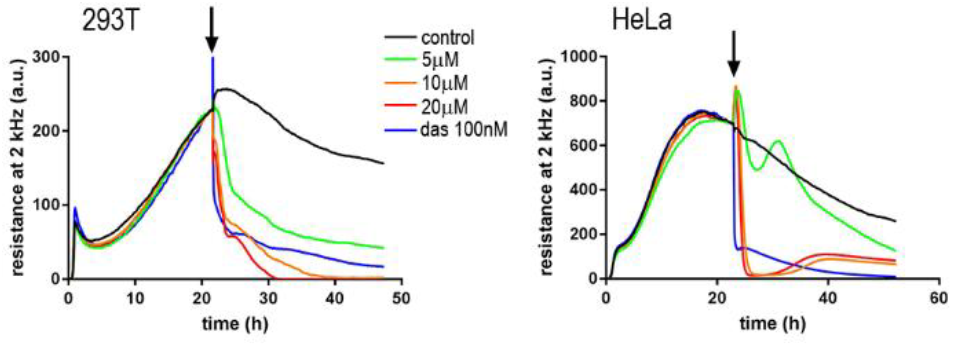
Decrease of the cell-surface contact area after NSC348884 addition. Resistance at 2 kHz was tracked during 293T (left) or HeLa (right) adhesion to the well bottom and after NSC348884 or dasatinib (das) addition. Times of the drug addition are marked by arrows. The curves represent mean values of triplicate/duplicate wells for NSC348884/das, respectively.

**Fig.14:**
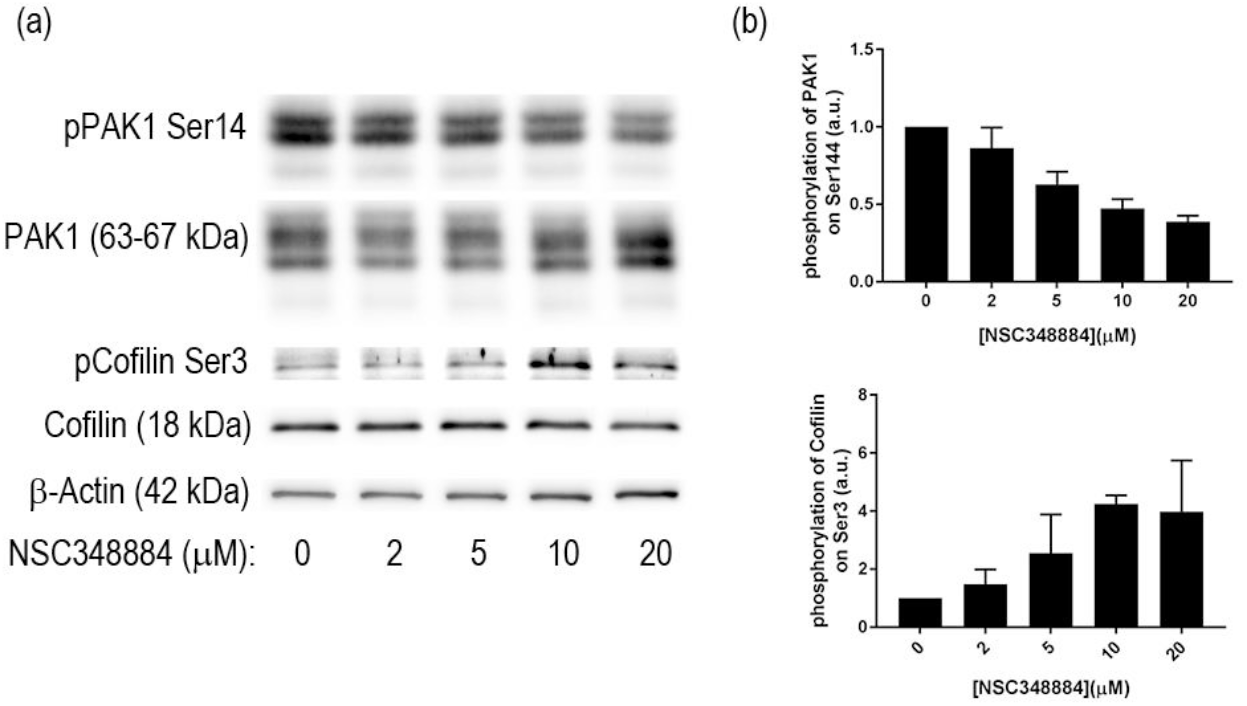
Expression and phosphorylation of adhesion-related protein kinase PAK1 and of the actin regulator Cofilin in 293T cells after 2h NSC348884 treatment. (a) Representative blots, (b) summary from 2 experiments. Error bars: ±SD

## Discussion

The N-terminal region of NPM is essential for its chaperone function as numerous proteins interact with NPM through this domain. Simultaneously, the domain is important for NPM oligomerization. Since AML-related NPM mutation does not affect its ability to form oligomers, NPM-interacting proteins become frequently mislocalized due to their interaction with aberrantly localized NPMmut. Targeting the NPM oligomerization offers a possibility to manipulate localization of the interacting partners. Although several alterations of NPM N-terminal domain were reported to disrupt NPM oligomerization *in vitro*, results thoroughly describing NPMwt and NPMmut interactions *in vivo* are missing. We have previously documented that C21 point mutations do not disrupt NPM oligomerization in live cells (35). Here we demonstrate that the cytoplasmic mislocalization of R_C21 and R_NPMwt in cells co-expressing G_NPMmut is very comparable (Fig. 1). This strongly indicates an existence of interaction between native NPM forms and the C21 mutants. Our results from native and semi-native electrophoreses allowed us to evaluate the interaction potential of C21 mutants depending on the substituted aminoacid (Fig. 2) *in vitro*, in agreement with the results of Prinos et al (60). Whereas C21F substitution significantly affected the stability of NPM oligomers at native conditions (Fig. 3 and S1), the effect of the C21A substitution was only detectable under reducing conditions. NPM oligomers were reported to consist of five NPM molecules (15), and formation of heterooligomers containing NPMwt and NPM mutants were found to be highly frequent (3, 7). We therefore suggest that the *in vivo* stability of NPM heterooligomers is permitted by a sufficient number of NPMwt molecules in the heterooligomeric complex. Under such conditions, the C21 point mutations do not have the potential to disrupt these complexes in living cells, although their stability is compromised.

In agreement with results of Enomoto et al (16), partial or complete deletion of NPM oligomerization domain (aa1-117) led to delocalization of the truncated protein from the nucleoli to the nucleoplasm (Fig. 4). However, even the NPM with completely deleted N-domain (Δ117) exhibited higher concentration in the nucleoli compared to the nucleoplasm. This is likely due to the fact, that nucleolar localization signal as well as nucleic acid binding domains remain unaffected by the deletion. We have found that oligomerization of the deletion mutants was completely abrogated and no interaction between two truncated NPM forms was detected (Figs 5, S2, S3). Nevertheless, immunoprecipitation revealed presence of both exo- and endogenous NPMwt in the G_Δ117 precipitates indicating that the truncated NPM yet participates in complexes, which are possibly too large to enter the native gel. The existence of mixed Δ117/NPMwt complexes is also supported by the FRET results monitoring the Δ117-NPMwt interaction in live cells (Fig. 6). Enhanced level of nucleolar proteins NCL and FBL co-precipitated with the deletion mutants suggests better accessibility of the NPM region responsible for binding of these proteins. Complexes containing NCL and/or FBL together with NPMwt and Δ117mutants thus represent a potential pool of proteins that can co-precipitate with the deletion mutants.

NSC348884 is declared to inhibit NPM oligomerization (61). We therefore analyzed its effect on various leukemia cell lines and on cells expressing fluorescently labeled NPM constructs. First, we tested cell viability and apoptotic signatures in order to determine the range of proper NSC348884 concentrations for the live-cell experiments. Concentrations required for a substantial viability decrease and caspase-3 fragmentation fell into the interval of 2-10μM for the majority of the cell lines (Fig. 7). Neither the OCI-AML3 cell line nor the primary cells of AML patients with NPM mutation displayed enhanced sensitivity to NSC348884 treatment. Interestingly, we noticed NSC34884-induced reduction of p14Arf in 293T and HeLa cells (data not shown). Unexpectedly, none of the tested cell lines nor primary cells from AML patients displayed any change in the NPM oligomerization upon treatment with efficient NSC348884 concentrations when investigated by the native and semi-native electrophoreses (Fig. 8). Similarly, oligomers containing fluorescently labeled NPMwt and NPMmut were not affected by the NSC348884 treatment (Fig. 10 and S4). These results were further verified by immunoprecipitation: both exogenous and endogenous NPM co-precipitated with both GFP- and RFP-labeled NPMwt and NPMmut, despite the presence of NSC348884 (Fig. 11). Also the fluorescence microscopy revealed sustained fraction of NPMwt in the cytoplasm of NSC348884 treated cells and FLIM-FRET imaging proved persisting interaction between fluorescently labeled NPM molecules upon the NSC348884 treatment (Fig. 12). Cells expressing two fluorescent variants of NCL were used as a control (Fig. S5). As expected, NCL molecules labeled with the eGFP donor and the mRFP1 acceptor on their N-termini did not exhibit any FRET, which was proved by a zero lifetime change upon the acceptor photobleaching. The result was independent of the NSC348884 treatment. Compared to NCL, the presence of FRET in the cells with fluorescently labeled NPM is clearly detectable both before and after the NSC348884 treatment. We conclude, that contrary to the published data (61), NSC348884 does not act as an oligomerization inhibitor and does not affect the stability of NPM oligomers under physiological conditions. This finding is extremely important in view of the fact that this drug has been recently reported to cause numerous cellular effects, which were ascribed, in accordance with its declared function, to disruption of NPM oligomerization (36, 56, 64).

During the live-cell experiments, we noticed apparent changes in cell adhesivity. The cell-surface contact area during the NSC348884 treatment was therefore monitored by Electrical Cell-Substrate Impedance Sensing (ECIS) (Fig.13). The rapid onset of changes in the ECIS signal indicated that the cell shrinkage and detachment was not a secondary effect accompanying apoptosis. As the course of the ECIS signal was similar to that induced by the inhibitor of p21-activated kinases, IPA-3 (27), we investigated also the level of activity and expression of PAK1 and of a known actin regulator, Cofilin (23) (Fig.14). The observed changes of both PAK1 and cofilin phosphorylation indicate that NSC348884 interferes with adhesion signaling. Further research is required to elucidate role of NSC348884 in this process and its potential for anticancer therapy.

## Conclusion

A set of complementary methods introduced here provides a reliable platform for investigation of NPM oligomerization and interaction network in both cell lysates and live cells. Our findings prove that point mutations in Cysteine 21 slightly modulate the oligomer stability but the NPM interaction potential is conserved. Deletion mutants lacking part of the NPM N-terminal domain completely lose their oligomerization ability, but they partially retain the interaction with NPMwt, possibly through enhanced interaction with other nucleolar proteins in complexes with NPMwt. We have shown that a proposed inhibitor of NPM oligomerization, NSC348884, does not affect NPM oligomer formation and stability in any of the examined leukemia cells. Moreover, the sensitivity to NSC348884 treatment is not potentiated by AML-associated NPM mutation. On the other hand, we uncovered an unknown effect of NSC48884 treatment on the cell-surface adhesion.

## Abbreviations

NPMwt: wild-type nucleophosmin
NPMmut: nucleophosmin with AML-associated mutation type A
C21: Cysteine 21 in nucleophosmin
C21A: substitution of C21 to Alanine
C21F: substitution of C21 to Phenylalanine
Δ25: nucleophosmin with deletion of the first 25 amino acids
Δ100: nucleophosmin with deletion of the first 100 amino acids
Δ117: nucleophosmin with deletion of the first 117 amino acids
G_: eGFP-labeled protein at the N-terminus
R_: mRFP1-labeled protein at the N-terminus

All authors declare no conflict of interest.

## Acknowledgements

The work was supported by the Czech Science Foundation (grant No 19-04099S), the Ministry of Health of the Czech Republic (project for conceptual development of the research organization No 00023736) and by the EU Operational Program OP VaVpI No. CZ.1.05/4.1.00/16.0340 provided by the Ministry of Education, CZ.

## Author contributions

BB, MŠ, PH, AH and KK conceived the experiments; all authors performed the research and analyzed data; MŠ and BB wrote the paper; BB, PH, KK and AH edited the manuscript.

## References

1. Albiero, E., D. Madeo, N. Bolli, I. Giaretta, E. D. Bona, M. F. Martelli, I. Nicoletti, F. Rodeghiero, and B. Falini. 2007. Identification and functional characterization of a cytoplasmic nucleophosmin leukaemic mutant generated by a novel exon-11 NPM1 mutation. Leukemia. 21: 1099-1103. doi: 2404597 [pii].

2. Balusu, R., W. Fiskus, R. Rao, D. G. Chong, S. Nalluri, U. Mudunuru, H. Ma, L. Chen, S. Venkannagari, K. Ha, S. Abhyankar, C. Williams, J. McGuirk, H. J. Khoury, C. Ustun, and K. N. Bhalla. 2011. Targeting levels or oligomerization of nucleophosmin 1 induces differentiation and loss of survival of human AML cells with mutant NPM1. Blood. 118: 3096-3106. doi: 10.1182/blood-2010-09-309674 [doi].

3. Bolli, N., M. F. De Marco, M. P. Martelli, B. Bigerna, A. Pucciarini, R. Rossi, R. Mannucci, N. Manes, V. Pettirossi, S. A. Pileri, I. Nicoletti, and B. Falini. 2009. A dose-dependent tug of war involving the NPM1 leukaemic mutant, nucleophosmin, and ARF. Leukemia. 23: 501-509. doi: 10.1038/leu.2008.326 [doi].

4. Borer, R. A., C. F. Lehner, H. M. Eppenberger, and E. A. Nigg. 1989. Major nucleolar proteins shuttle between nucleus and cytoplasm. Cell. 56: 379–390.

5. Borrow, J., S. A. Dyer, S. Akiki, and M. J. Griffiths. 2019. Molecular roulette: nucleophosmin mutations in AML are orchestrated through N-nucleotide addition by TdT. Blood. 134: 2291-2303. doi: 10.1182/blood.2019001240. https://ashpublications.org/blood/article/134/25/2291/421222/Molecular-roulette-nucleophosmin-mutations-in-AML.

6. Brodska, B., A. Holoubek, P. Otevrelova, and K. Kuzelova. 2016. Low-Dose Actinomycin-D Induces Redistribution of Wild-Type and Mutated Nucleophosmin Followed by Cell Death in Leukemic Cells. J. Cell. Biochem. 117: 1319-1329. doi: 10.1002/jcb.25420 [doi].

7. Brodska, B., M. Kracmarova, A. Holoubek, and K. Kuzelova. 2017. Localization of AML-related nucleophosmin mutant depends on its subtype and is highly affected by its interaction with wild-type NPM. PLoS One. 12: e0175175. doi: 10.1371/journal.pone.0175175 [doi].

8. Campregher, P. V., W. de Oliveira Pereira, B. Lisboa, R. Puga, E. R. Deolinda, R. Helman, L. C. Marti, J. C. Guerra, K. N. Manola, R. C. Petroni, A. M. Bezerra, F. F. Costa, N. Hamerschlak, and de Souza Santos, F P. 2016. A novel mechanism of NPM1 cytoplasmic localization in acute myeloid leukemia: the recurrent gene fusion NPM1-HAUS1. Haematologica. 101: 287. doi: 10.3324/haematol.2015.137364 [doi].

9. Chou, S., B. Ko, J. Chiou, Y. Hsu, M. Tsai, Y. Chiu, I. -. Yu, S. Lin, H. Hou, Y. Kuo, H. Lin, M. Wu, W. Chou, and H. Tien. 2012. A knock-in Npm1 mutation in mice results in myeloproliferation and implies a perturbation in hematopoietic microenvironment. PLoS ONE. 7: e49769. doi: 10.1371/journal.pone.0049769.

10. Chung, E. Y., Y. Mai, U. A. Shah, Y. Wei, E. Ishida, K. Kataoka, X. Ren, K. Pradhan, B. Bartholdy, X. Wei, Y. Zou, J. Zhang, S. Ogawa, U. Steidl, X. Zang, A. Verma, M. Janakiram, and B. H. Ye. 2019. PAK Kinase Inhibition Has Therapeutic Activity in Novel Preclinical Models of Adult T-Cell Leukemia/Lymphoma. Clin. Cancer Res. 25: 3589-3601. doi: 10.1158/1078-0432.CCR-18-3033.

11. Colombo, E., J. C. Marine, D. Danovi, B. Falini, and P. G. Pelicci. 2002. Nucleophosmin regulates the stability and transcriptional activity of p53. Nat. Cell Biol. 4: 529-533. doi: 10.1038/ncb814.

12. Cordell, J. L., K. A. Pulford, B. Bigerna, G. Roncador, A. Banham, E. Colombo, P. G. Pelicci, D. Y. Mason, and B. Falini. 1999. Detection of normal and chimeric nucleophosmin in human cells. Blood. 93: 632–642.

13. den Besten, W., M. L. Kuo, R. T. Williams, and C. J. Sherr. 2005. Myeloid leukemia-associated nucleophosmin mutants perturb p53-dependent and independent activities of the Arf tumor suppressor protein. Cell. Cycle. 4: 1593-1598. doi: 2174 [pii].

14. Dohner, K., R. F. Schlenk, M. Habdank, C. Scholl, F. G. Rucker, A. Corbacioglu, L. Bullinger, S. Frohling, and H. Dohner. 2005. Mutant nucleophosmin (NPM1) predicts favorable prognosis in younger adults with acute myeloid leukemia and normal cytogenetics: interaction with other gene mutations. Blood. 106: 3740-3746. doi: 2005-05-2164 [pii].

15. Dutta, S., I. V. Akey, C. Dingwall, K. L. Hartman, T. Laue, R. T. Nolte, J. F. Head, and C. W. Akey. 2001. The crystal structure of nucleoplasmin-core: implications for histone binding and nucleosome assembly. Mol. Cell. 8: 841–853.

16. Enomoto, T., M. S. Lindstrom, A. Jin, H. Ke, and Y. Zhang. 2006. Essential role of the B23/NPM core domain in regulating ARF binding and B23 stability. J. Biol. Chem. 281: 18463-18472. doi: M602788200 [pii].

17. Falini, B., N. Bolli, A. Liso, M. P. Martelli, R. Mannucci, S. Pileri, and I. Nicoletti. 2009. Altered nucleophosmin transport in acute myeloid leukaemia with mutated NPM1: molecular basis and clinical implications. Leukemia. 23: 1731-1743. doi: 10.1038/leu.2009.124 [doi].

18. Falini, B., N. Bolli, J. Shan, M. P. Martelli, A. Liso, A. Pucciarini, B. Bigerna, L. Pasqualucci, R. Mannucci, R. Rosati, P. Gorello, D. Diverio, G. Roti, E. Tiacci, G. Cazzaniga, A. Biondi, S. Schnittger, T. Haferlach, W. Hiddemann, M. F. Martelli, W. Gu, C. Mecucci, and I. Nicoletti. 2006. Both carboxy-terminus NES motif and mutated tryptophan(s) are crucial for aberrant nuclear export of nucleophosmin leukemic mutants in NPMc+ AML. Blood. 107: 4514-4523. doi: 2005-11-4745 [pii].

19. Falini, B., M. P. Martelli, N. Bolli, R. Bonasso, E. Ghia, M. T. Pallotta, D. Diverio, I. Nicoletti, R. Pacini, A. Tabarrini, B. V. Galletti, R. Mannucci, G. Roti, R. Rosati, G. Specchia, A. Liso, E. Tiacci, M. Alcalay, L. Luzi, S. Volorio, L. Bernard, A. Guarini, S. Amadori, F. Mandelli, F. Pane, F. Lo-Coco, G. Saglio, P. G. Pelicci, M. F. Martelli, and C. Mecucci. 2006. Immunohistochemistry predicts nucleophosmin (NPM) mutations in acute myeloid leukemia. Blood. 108: 1999-2005. doi: blood-2006-03-007013 [pii].

20. Falini, B., C. Mecucci, E. Tiacci, M. Alcalay, R. Rosati, L. Pasqualucci, R. La Starza, D. Diverio, E. Colombo, A. Santucci, B. Bigerna, R. Pacini, A. Pucciarini, A. Liso, M. Vignetti, P. Fazi, N. Meani, V. Pettirossi, G. Saglio, F. Mandelli, F. Lo-Coco, P. G. Pelicci, M. F. Martelli, and GIMEMA Acute Leukemia Working Party. 2005. Cytoplasmic nucleophosmin in acute myelogenous leukemia with a normal karyotype. N. Engl. J. Med. 352: 254-266. doi: 352/3/254 [pii].

21. Falini, B., I. Nicoletti, M. F. Martelli, and C. Mecucci. 2007. Acute myeloid leukemia carrying cytoplasmic/mutated nucleophosmin (NPMc+ AML): biologic and clinical features. Blood. 109: 874-885. doi: blood-2006-07-012252 [pii].

22. Federici, L., and B. Falini. 2013. Nucleophosmin mutations in acute myeloid leukemia: a tale of protein unfolding and mislocalization. Protein Sci. 22: 545-556. doi: 10.1002/pro.2240 [doi].

23. Flis, S., E. Bratek, T. Chojnacki, M. Piskorek, and T. Skorski. 2019. Simultaneous Inhibition of BCR-ABL1 Tyrosine Kinase and PAK1/2 Serine/Threonine Kinase Exerts Synergistic Effect against Chronic Myeloid Leukemia Cells. Cancers (Basel). 11:. doi: 10.3390/cancers11101544.

24. Frehlick, L. J., J. M. Eirín-López, and J. Ausió. 2007. New insights into the nucleophosmin/nucleoplasmin family of nuclear chaperones. Bioessays. 29: 49-59. doi: 10.1002/bies.20512.

25. Garzon, R., M. Savona, R. Baz, M. Andreeff, N. Gabrail, M. Gutierrez, L. Savoie, P. Mau- Sorensen, N. Wagner-Johnston, K. Yee, T. J. Unger, J. Saint-Martin, R. Carlson, T. Rashal, T. Kashyap, B. Klebanov, S. Shacham, M. Kauffman, and R. Stone. 2017. A phase 1 clinical trial of single-agent selinexor in acute myeloid leukemia. Blood. 129: 3165-3174. doi: 10.1182/blood-2016-11-750158.

26. Gorello, P., G. Cazzaniga, F. Alberti, M. G. Dell’Oro, E. Gottardi, G. Specchia, G. Roti, R. Rosati, M. F. Martelli, D. Diverio, F. Lo Coco, A. Biondi, G. Saglio, C. Mecucci, and B. Falini. 2006. Quantitative assessment of minimal residual disease in acute myeloid leukemia carrying nucleophosmin (NPM1) gene mutations. Leukemia. 20: 1103-1108. doi: 10.1038/sj.leu.2404149.

27. Grebeňová, D., A. Holoubek, P. Röselová, A. Obr, B. Brodská, and K. Kuželová. 2019. PAK1, PAK1Δ15, and PAK2: similarities, differences and mutual interactions. Sci Rep. 9: 17171. doi: 10.1038/s41598-019-53665-6.

28. Grisendi, S., R. Bernardi, M. Rossi, K. Cheng, L. Khandker, K. Manova, and P. P. Pandolfi. 2005. Role of nucleophosmin in embryonic development and tumorigenesis. Nature. 437: 147-153. doi: nature03915 [pii].

29. Grisendi, S., C. Mecucci, B. Falini, and P. P. Pandolfi. 2006. Nucleophosmin and cancer. Nat. Rev. Cancer. 6: 493-505. doi: 10.1038/nrc1885.

30. Gu, X., Q. Ebrahem, R. Z. Mahfouz, M. Hasipek, F. Enane, T. Radivoyevitch, N. Rapin, B. Przychodzen, Z. Hu, R. Balusu, C. V. Cotta, D. Wald, C. Argueta, Y. Landesman, M. P. Martelli, B. Falini, H. Carraway, B. T. Porse, J. Maciejewski, B. K. Jha, and Y. Saunthararajah. 2018. Leukemogenic nucleophosmin mutation disrupts the transcription factor hub that regulates granulomonocytic fates. J. Clin. Invest. 128: 4260-4279. doi: 10.1172/JCI97117.

31. Heiblig, M., H. Labussière-Wallet, F. E. Nicolini, M. Michallet, S. Hayette, P. Sujobert, A. Plesa, M. Balsat, E. Paubelle, F. Barraco, I. Tigaud, S. Ducastelle, E. Wattel, G. Salles, and X. Thomas. 2019. Prognostic Value of Genetic Alterations in Elderly Patients with Acute Myeloid Leukemia: A Single Institution Experience. Cancers (Basel). 11:. doi: 10.3390/cancers11040570.

32. Herman, P., A. Holoubek, and B. Brodska. 2019. Lifetime-based photoconversion of EGFP as a tool for FLIM. Biochim Biophys Acta Gen Subj. 1863: 266-277. doi: 10.1016/j.bbagen.2018.10.016.

33. Herrera, J. E., J. J. Correia, A. E. Jones, and M. O. Olson. 1996. Sedimentation analyses of the salt- and divalent metal ion-induced oligomerization of nucleolar protein B23. Biochemistry. 35: 2668–2673. doi: 10.1021/bi9523320 [doi].

34. Herrera, J. E., R. Savkur, and M. O. Olson. 1995. The ribonuclease activity of nucleolar protein B23. Nucleic Acids Res. 23: 3974-3979. doi: 5a0309 [pii].

35. Holoubek, A., P. Herman, J. Sýkora, B. Brodská, J. Humpolíčková, M. Kráčmarová, D. Gášková, M. Hof, and K. Kuželová. 2018. Monitoring of nucleophosmin oligomerization in live cells. Methods Appl Fluoresc. 6: 035016. doi: 10.1088/2050-6120/aaccb9.

36. Hu, W., Y. Liang, J. Luo, X. Gu, Z. Chen, T. Fu, Y. Zhu, S. Lin, H. Diao, B. Jia, and Z. Yang. 2019. Nucleolar stress regulation of endometrial receptivity in mouse models and human cell lines. Cell Death Dis. 10: 831. doi: 10.1038/s41419-019-2071-6.

37. Huang, M., D. Thomas, M. X. Li, W. Feng, S. M. Chan, R. Majeti, and B. S. Mitchell. 2013. Role of cysteine 288 in nucleophosmin cytoplasmic mutations: sensitization to toxicity induced by arsenic trioxide and bortezomib. Leukemia. 27: 1970-1980. doi: 10.1038/leu.2013.222 [doi].

38. Itahana, K., K. P. Bhat, A. Jin, Y. Itahana, D. Hawke, R. Kobayashi, and Y. Zhang. 2003. Tumor suppressor ARF degrades B23, a nucleolar protein involved in ribosome biogenesis and cell proliferation. Mol. Cell. 12: 1151–1164.

39. Konoplev, S., X. Huang, H. A. Drabkin, H. Koeppen, D. Jones, H. M. Kantarjian, G. Garcia- Manero, W. Chen, L. J. Medeiros, and C. E. Bueso-Ramos. 2009. Cytoplasmic localization of nucleophosmin in bone marrow blasts of acute myeloid leukemia patients is not completely concordant with NPM1 mutation and is not predictive of prognosis. Cancer. 115: 4737-4744. doi: 10.1002/cncr.24543 [doi].

40. Kuzelova, K., B. Brodska, O. Fuchs, M. Dobrovolna, P. Soukup, and P. Cetkovsky. 2015. Altered HLA Class I Profile Associated with Type A/D Nucleophosmin Mutation Points to Possible Anti-Nucleophosmin Immune Response in Acute Myeloid Leukemia. PLoS One. 10: e0127637. doi: 10.1371/journal.pone.0127637 [doi].

41. Kuželová, K., B. Brodská, J. Schetelig, C. Röllig, Z. Ráčil, J. S. Walz, G. Helbig, O. Fuchs, M. Vraná, P. Pecherková, C. Šálek, and J. Mayer. 2018. Association of HLA class I type with prevalence and outcome of patients with acute myeloid leukemia and mutated nucleophosmin. PLoS ONE. 13: e0204290. doi: 10.1371/journal.pone.0204290.

42. Lee, H. H., H. S. Kim, J. Y. Kang, B. I. Lee, J. Y. Ha, H. J. Yoon, S. O. Lim, G. Jung, and S. W. Suh. 2007. Crystal structure of human nucleophosmin-core reveals plasticity of the pentamer-pentamer interface. Proteins. 69: 672-678. doi: 10.1002/prot.21504.

43. Li, J., X. Zhang, D. P. Sejas, G. C. Bagby, and Q. Pang. 2004. Hypoxia-induced nucleophosmin protects cell death through inhibition of p53. J. Biol. Chem. 279: 41275-41279. doi: 10.1074/jbc.C400297200.

44. Li, Y. P., R. K. Busch, B. C. Valdez, and H. Busch. 1996. C23 interacts with B23, a putative nucleolar-localization-signal-binding protein. Eur. J. Biochem. 237: 153–158.

45. Li, Z., D. Boone, and S. R. Hann. 2008. Nucleophosmin interacts directly with c-Myc and controls c-Myc-induced hyperproliferation and transformation. Proc. Natl. Acad. Sci. U. S. A. 105: 18794-18799. doi: 10.1073/pnas.0806879105 [doi].

46. Lindström, M. S. 2011. NPM1/B23: A Multifunctional Chaperone in Ribosome Biogenesis and Chromatin Remodeling. Biochem Res Int. 2011: 195209. doi: 10.1155/2011/195209.

47. Macville, M., E. Schrock, H. Padilla-Nash, C. Keck, B. M. Ghadimi, D. Zimonjic, N. Popescu, and T. Ried. 1999. Comprehensive and definitive molecular cytogenetic characterization of HeLa cells by spectral karyotyping. Cancer Res. 59: 141–150.

48. Mariano, A. R., E. Colombo, L. Luzi, P. Martinelli, S. Volorio, L. Bernard, N. Meani, R. Bergomas, M. Alcalay, and P. G. Pelicci. 2006. Cytoplasmic localization of NPM in myeloid leukemias is dictated by gain-of-function mutations that create a functional nuclear export signal. Oncogene. 25: 4376–4380. doi: 10.1038/sj.onc.1209453.

49. Meani, N., and M. Alcalay. 2009. Role of nucleophosmin in acute myeloid leukemia. Expert Rev. Anticancer Ther. 9: 1283-1294. doi: 10.1586/era.09.84 [doi].

50. Morris, S. W., M. N. Kirstein, M. B. Valentine, K. G. Dittmer, D. N. Shapiro, D. L. Saltman, and A. T. Look. 1994. Fusion of a kinase gene, ALK, to a nucleolar protein gene, NPM, in non-Hodgkin’s lymphoma. Science. 263: 1281–1284.

51. Namboodiri, V. M. H., S. Dutta, I. V. Akey, J. F. Head, and C. W. Akey. 2003. The crystal structure of Drosophila NLP-core provides insight into pentamer formation and histone binding. Structure. 11: 175–186.

52. Okuda, M. 2002. The role of nucleophosmin in centrosome duplication. Oncogene. 21: 6170-6174. doi: 10.1038/sj.onc.1205708.

53. Okuda, M., H. F. Horn, P. Tarapore, Y. Tokuyama, A. G. Smulian, P. K. Chan, E. S. Knudsen, I. A. Hofmann, J. D. Snyder, K. E. Bove, and K. Fukasawa. 2000. Nucleophosmin/B23 is a target of CDK2/cyclin E in centrosome duplication. Cell. 103: 127-140. doi: S0092-8674(00)00093-3 [pii].

54. Okuwaki, M., K. Matsumoto, M. Tsujimoto, and K. Nagata. 2001. Function of nucleophosmin/B23, a nucleolar acidic protein, as a histone chaperone. FEBS Lett. 506: 272-276. doi: S0014-5793(01)02939-8 [pii].

55. Patting, M. 2008. Evaluation of Time-Resolved Fluorescence Data: Typical Methods and Problems, p. 233-258. In Anonymous Standardization and Quality Assurance in Fluorescence Measurements Ivol. 5. Springer Berlin Heidelberg, Berlin, Heidelberg.

56. Phi, J. H., C. Sun, S. Lee, S. Lee, I. Park, S. A. Choi, S. Park, J. Y. Lee, K. Wang, S. Kim, H. Yun, and C. Park. 2019. NPM1 as a potential therapeutic target for atypical teratoid/rhabdoid tumors. BMC Cancer. 19: 848. doi: 10.1186/s12885-019-6044-z.

57. Pitiot, A. S., I. Santamaria, O. Garcia-Suarez, I. Centeno, A. Astudillo, C. Rayon, and M. Balbin. 2007. A new type of NPM1 gene mutation in AML leading to a C-terminal truncated protein. Leukemia. 21: 1564-1566. doi: 2404679 [pii].

58. Poletto M, Lirussi L, Wilson DM,Tell G. (2014) Nucleophosmin modulates stability, activity, and nucleolar accumulation of base excision repair proteins. Molecular Biology of the Cell 25(10):1641-1652. doi: 10.1091/mbc.e13-12-0717. https://www.ncbi.nlm.nih.gov/pmc/articles/PMC4019495/.

59. Prassek, V. V., M. Rothenberg-Thurley, M. C. Sauerland, T. Herold, H. Janke, B. Ksienzyk, N. P. Konstandin, D. Goerlich, U. Krug, A. Faldum, W. E. Berdel, B. Wörmann, J. Braess, S. Schneider, M. Subklewe, S. K. Bohlander, W. Hiddemann, K. Spiekermann, and K. H. Metzeler. 2018. Genetics of acute myeloid leukemia in the elderly: mutation spectrum and clinical impact in intensively treated patients aged 75 years or older. Haematologica. 103: 1853-1861. doi: 10.3324/haematol.2018.191536.

60. Prinos, P., M. C. Lacoste, J. Wong, A. M. Bonneau, and E. Georges. 2011. Mutation of cysteine 21 inhibits nucleophosmin/B23 oligomerization and chaperone activity. Int. J. Biochem. Mol. Biol. 2: 24–30.

61. Qi, W., K. Shakalya, A. Stejskal, A. Goldman, S. Beeck, L. Cooke, and D. Mahadevan. 2008. NSC348884, a nucleophosmin inhibitor disrupts oligomer formation and induces apoptosis in human cancer cells. Oncogene. 27: 4210-4220. doi: 10.1038/onc.2008.54 [doi].

62. Ranganathan, P., T. Kashyap, X. Yu, X. Meng, T. Lai, B. McNeil, B. Bhatnagar, S. Shacham, M. Kauffman, A. M. Dorrance, W. Blum, D. Sampath, Y. Landesman, and R. Garzon. 2016. XPO1 Inhibition using Selinexor Synergizes with Chemotherapy in Acute Myeloid Leukemia by Targeting DNA Repair and Restoring Topoisomerase IIα to the Nucleus. Clin. Cancer Res. 22: 6142-6152. doi: 10.1158/1078-0432.CCR-15-2885.

63. Redner, R. L., E. A. Rush, S. Faas, W. A. Rudert, and S. J. Corey. 1996. The t(5;17) variant of acute promyelocytic leukemia expresses a nucleophosmin-retinoic acid receptor fusion. Blood. 87: 882–886.

64. Reichert, F., and S. Rotshenker. 2019. Galectin-3 (MAC-2) Controls Microglia Phenotype Whether Amoeboid and Phagocytic or Branched and Non-phagocytic by Regulating the Cytoskeleton. Front Cell Neurosci. 13: 90. doi: 10.3389/fncel.2019.00090.

65. Röselová, P., A. Obr, A. Holoubek, D. Grebeňová, and K. Kuželová. 2018. Adhesion structures in leukemia cells and their regulation by Src family kinases. Cell Adh Migr. 12: 286-298. doi: 10.1080/19336918.2017.1344796.

66. Šašinková, M., A. Holoubek, P. Otevřelová, K. Kuželová, and B. Brodská. 2018. AML-associated mutation of nucleophosmin compromises its interaction with nucleolin. Int. J. Biochem. Cell Biol. 103: 65-73. doi: 10.1016/j.biocel.2018.08.008.

67. Savkur, R. S., and M. O. Olson. 1998. Preferential cleavage in pre-ribosomal RNA byprotein B23 endoribonuclease. Nucleic Acids Res. 26: 4508–4515.

68. Schindelin, J., I. Arganda-Carreras, E. Frise, V. Kaynig, M. Longair, T. Pietzsch, S. Preibisch, C. Rueden, S. Saalfeld, B. Schmid, J. Tinevez, D. J. White, V. Hartenstein, K. Eliceiri, P. Tomancak, and A. Cardona. 2012. Fiji: an open-source platform for biological-image analysis. Nat. Methods. 9: 676-682. doi: 10.1038/nmeth.2019.

69. Schmidt-Zachmann, M. S., B. Hugle-Dorr, and W. W. Franke. 1987. A constitutive nucleolar protein identified as a member of the nucleoplasmin family. Embo J. 6: 1881–1890.

70. Schnittger, S., C. Schoch, W. Kern, C. Mecucci, C. Tschulik, M. F. Martelli, T. Haferlach, W. Hiddemann, and B. Falini. 2005. Nucleophosmin gene mutations are predictors of favorable prognosis in acute myelogenous leukemia with a normal karyotype. Blood. 106: 3733-3739. doi: 10.1182/blood-2005-06-2248.

71. Takemura, M., N. Ohta, Y. Furuichi, T. Takahashi, S. Yoshida, M. O. Olson, and H. Umekawa. 1994. Stimulation of calf thymus DNA polymerase alpha activity by nucleolar protein B23. Biochem. Biophys. Res. Commun. 199: 46-51. doi: 10.1006/bbrc.1994.1191.

72. Takemura, M., K. Sato, M. Nishio, T. Akiyama, H. Umekawa, and S. Yoshida. 1999. Nucleolar protein B23.1 binds to retinoblastoma protein and synergistically stimulates DNA polymerase alpha activity. J. Biochem. 125: 904–909.

73. Verhaak, R. G., C. S. Goudswaard, W. van Putten, M. A. Bijl, M. A. Sanders, W. Hugens, A. G. Uitterlinden, C. A. Erpelinck, R. Delwel, B. Lowenberg, and P. J. Valk. 2005. Mutations in nucleophosmin (NPM1) in acute myeloid leukemia (AML): association with other gene abnormalities and previously established gene expression signatures and their favorable prognostic significance. Blood. 106: 3747-3754. doi: 2005-05-2168 [pii].

74. Wang, W., A. Budhu, M. Forgues, and X. W. Wang. 2005. Temporal and spatial control of nucleophosmin by the Ran-Crm1 complex in centrosome duplication. Nat. Cell Biol. 7: 823-830. doi: 10.1038/ncb1282.

75. Wu, M. H., J. H. Chang, and B. Y. M. Yung. 2002. Resistance to UV-induced cell-killing in nucleophosmin/B23 over-expressed NIH 3T3 fibroblasts: enhancement of DNA repair and up-regulation of PCNA in association with nucleophosmin/B23 over-expression. Carcinogenesis. 23: 93–100.

76. Yoneda-Kato, N., A. T. Look, M. N. Kirstein, M. B. Valentine, S. C. Raimondi, K. J. Cohen, A. J. Carroll, and S. W. Morris. 1996. The t(3;5)(q25.1;q34) of myelodysplastic syndrome and acute myeloid leukemia produces a novel fusion gene, NPM-MLF1. Oncogene. 12: 265–275.

77. Ziv, O., A. Zeisel, N. Mirlas-Neisberg, U. Swain, R. Nevo, N. Ben-Chetrit, M. P. Martelli, R. Rossi, S. Schiesser, C. E. Canman, T. Carell, N. E. Geacintov, B. Falini, E. Domany, and Z. Livneh. 2014. Identification of novel DNA-damage tolerance genes reveals regulation of translesion DNA synthesis by nucleophosmin. Nat Commun. 5:5437. doi: 10.1038/ncomms6437.

